# In-cell structure and snapshots of *copia* retrotransposons in intact tissue by cryo-electron tomography

**DOI:** 10.1101/2024.02.21.581285

**Authors:** Sven Klumpe, Kirsten A. Senti, Florian Beck, Jenny Sachweh, Bernhard Hampoelz, Paolo Ronchi, Assa Yeroslaviz, John A.G. Briggs, Julius Brennecke, Martin Beck, Jürgen M. Plitzko

## Abstract

Long terminal repeat (LTR) retrotransposons belong to the transposable elements (TE), autonomously replicating genetic elements that integrate into the host’s genome. LTR retrotransposons represent a major component of genomes across the tree of life; some derived sequences have even been domesticated by the host to perform cellular functions in essential processes such as development. Among animals, *Drosophila melanogaster* serves as an important model organism for TE research, harboring several LTR retrotransposons, including the Ty1-*copia* family, which is evolutionarily related to retroviruses and forms virus-like particles (VLPs). The architectural organization of *copia* VLPs *in situ* has remained unknown. In this study, we use cryo-FIB milling and lift-out approaches to visualize *copia* VLPs in isolated ovarian cells and intact egg chambers and resolve the *in situ copia* capsid structure to 7.7 Å resolution by cryo-ET. While cytosolic *copia* VLPs vary in size, nuclear VLPs are homogenous and form densely packed clusters, supporting a model in which nuclear import acts as a size selector. By analyzing flies deficient in the TE-suppressing PIWI-piRNA pathway, we observe a change in *copia* localization from cytosolic to nuclear during spermatogenesis in testes. Our findings provide insights into the cellular structural biology of an active LTR retrotransposon and shed light on the replication cycle of *copia* in the context of host gametogenesis.

## Introduction

Transposable elements (TEs) are a diverse group of mobile genetic elements that can autonomously replicate and insert themselves into new genomic locations [1, 2]. TEs are widespread in organisms across the tree of life, frequently constituting a substantial proportion of their host genomes. In mammals, for instance, TEs account for approximately 25-75% of the genome, with humans reaching ∼50% [3–5], and in some plant species, TEs can comprise up to 85% of the genome [6]. A subclass of TEs, long terminal repeat (LTR) retrotransposons, transpose via an RNA intermediate and are evolutionarily related to retroviruses [7]. Their genomic architecture is characterized by two LTRs containing regulatory sequences and flanking the two major open reading frames (ORFs), *gag* encoding the structural proteins capsid (CA) and nucleocapsid (NC), and *pol* encoding the enzymes protease (PR), integrase (IN) and reverse transcriptase (RT). The Ty1-*copia* LTR retrotransposons are classified as the virus family of Pseudoviridae and are grouped in the order Ortervirales that also encompasses Retroviridae [8]. The members of the Ty1-*copia* group differ from other LTR retroelements in their genomic structure: (1) they encode a single open reading frame, (2) they undergo alternative splicing in animals [9] and plants [10], and (3) they have a reverse order of the IN and RT coding sequences [11].

Ty1-*copia* elements and LTR retrotransposons in general undergo a replication cycle similar to that of retroviruses, involving transcription from their genomic copies, nuclear export of the genomic RNA, translation of the encoded ORFs in the cytoplasm, capsid formation, and finally, reverse transcription of their RNA genome, nuclear import and integration of a new copy into the host genome. The events from reverse transcription to integration are not fully resolved, and the exact order of the contributing molecular processes may differ between elements. A crucial question in the replication cycle of retroviruses and LTR retrotransposons is how they enter the nucleus to enable integration into the genome. Some retroviruses (e.g. murine leukemia virus, MuLV) require nuclear breakdown to gain access to host chromosomes [12, 13]. In the case of human immunodeficiency virus 1 (HIV-1), the mature cone-shaped nucleocapsid containing the partially reverse transcribed HIV-1 genome can translocate into the nucleus as an intact entity through nuclear pore complexes [14]. Hereby, they directly interact with central channel nucleoporins [15, 16] and, thus, do not require nuclear transport receptors for their nuclear entry, despite their extraordinary size.

In animals, TEs are linked to the germline as their evolutionary success depends on vertical transmission [17]. It is therefore not surprising that LTR retrotransposons are expressed predominantly in the host gonads where they hijack the cellular machinery for their own replication and propagation within the host’s germline genome. To safeguard the genomic integrity of the germline lineage, animals have evolved intricate defense mechanisms targeted against TEs. A central pillar of this defense repertoire is the PIWI-interacting RNA (piRNA) pathway, which mediates TE silencing, and is centered around Argonaute proteins belonging to the PIWI clade and their associated 22-32 nucleotide long piRNAs [18, 19]. Derived from TE-sequences, piRNAs guide PIWI-clade proteins to complementary TE transcripts instigating both transcriptional and post-transcriptional TE silencing. As the piRNA pathway silences most TEs simultaneously and as the experimental ablation of specific piRNA populations is often not possible, dissecting the replication cycle of an individual TE presents a formidable challenge.

Here, we focus on the fruit fly *Drosophila melanogaster*, a key model organism for studying TEs and their co-evolution with a host organism [20], to investigate the replication cycle of the *copia* LTR retrotransposon. We find that *copia* stands out among LTR retrotransposons in that it is expressed at high levels in *Drosophila* ovaries with an intact piRNA pathway, specifically in somatic follicle cells that surround the germline in the egg chamber (Fig. 1A). Employing cryo-electron tomography (cryo-ET) on cryo-focused ion beam (FIB) milled lamellae [21, 22], we visualize *copia* virus-like particles (VLPs) within their native host cells and use subtomogram analysis to resolve its *in situ* capsid *s*tructure to 7.7 Å resolution, allowing us to describe key architectural features of its capsid. Using cryo-lift-out techniques [23–26], we find *copia* capsids both inside and outside the nucleus in the ovarian follicle cells of the intact *D. melanogaster* egg chamber. In several instances, cytoplasmic *copia* capsids are in the direct vicinity of the nuclear pore complex (NPC) and engage with its central channel. By employing piRNA pathway genetics in combination with fluorescent RNA *in situ* hybridization (RNA FISH) and immuno-fluorescence experiments, we elucidate the strict repression and nuclear translocation of *copia* in germline cells of the developing testes. Using a multimodal approach including novel techniques for cryo-ET in cells and tissues [27], our work demonstrates the *in situ* characterization of an active LTR retrotransposon in its relevant host environment.

**Figure 1:**
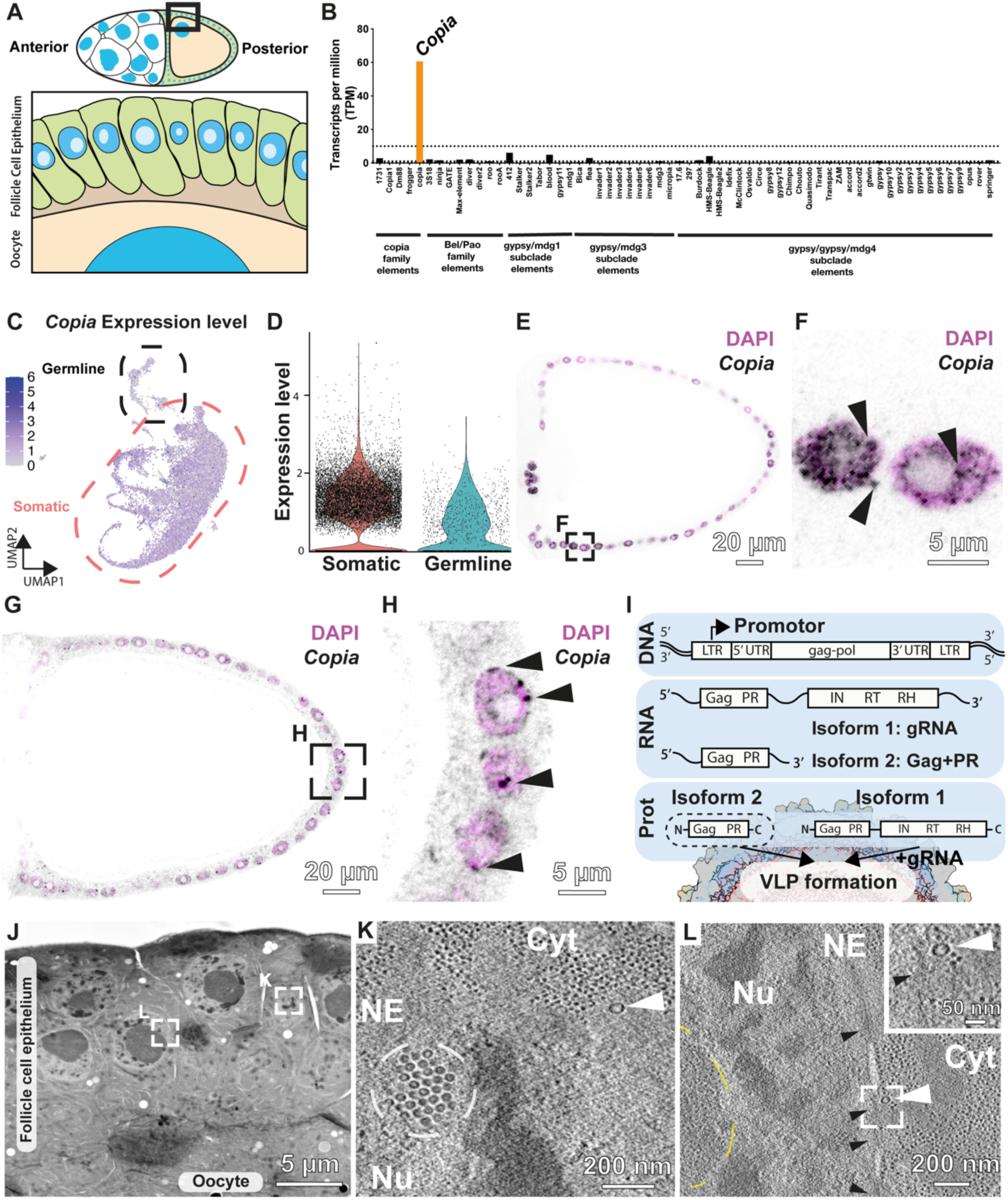
*Copia* is an abundant retrotransposon that forms VLPs in the ovarian follicle cells. **A:** Schematic of a stage 10 egg chamber. Shown are somatic follicle cells (green) and the germline that encompasses nurse cells (white) and the oocyte (beige) with nuclei shown in blue and nucleoli shown in light blue. In the zoom on the follicle cell epithelium, the area between follicle cell and oocyte (brown) represents the vitelline membrane. **B:** Plot showing expression levels of all LTR elements present in the *D. melanogaster* genome using polyA-RNA-seq from bulk ovaries of the control RNAi genotype (*tj>arrestin2/GD*) in transcripts per million (TPM). *Copia* is highlighted in orange. **C:** scRNA-seq analysis of *copia* in the *D. melanogaster* of *w^-^* ovary. Shown is the normalized expression in arbitrary units. **D:** Violin plot of data from C showing *copia* expression levels in the somatic cells versus germline cells. **E,F:** Color-inverted confocal images showing *copia* smFISH (black) and DAPI (magenta) of a control knockdown genotype (*tj>arrestin2/GD)* stage 10 egg chamber. *Copia* transcript is detected in follicle cells while absent in the oocyte as well as nurse cells (compare Fig. S2A). **F:** Magnified panel from E. *Copia* transcript levels are high in follicle cell nuclei and their periphery (black arrowheads) **G, H:** Confocal image of anti-*copia* Gag staining (black) and DAPI (magenta) of a stage 10 egg chamber of a control knockdown (*tj>white sh*). **H:** Magnified panel from G. Black arrowheads show *copia* signal accumulation in the cytoplasm, nucleoplasm and at the nuclear envelope. **I**: Schematic representation of *copia* transcription and translation. Alternative splicing of the *copia* transcript leads to the RNA genome encoding the full-length polypeptide and an isoform that encodes Gag and PR. The gRNA and the two proteins isoforms form *copia* VLPs. **J**: Overview image of a high-pressure frozen, freeze-substituted and resin-embedded TEM section from a stage 10 egg chamber of *tj>Sec31*::*mCherry* ovary showing the follicle cell epithelium. White rectangles indicate tomogram positions. **K-L**: Tomographic slices of the nuclear periphery of the follicle cell nucleus showing VLP clusters in the nucleus (white dashed circle), cytoplasmic particles (white arrowhead), nuclear pores (black arrowheads), and the nucleolus (yellow dashed circle). **L inset**: a VLP in proximity of a nuclear pore.

## Results

### Abundant virus-like particles of the *copia* LTR retrotransposon in ovaries

Consistent with the general silencing capacity of the piRNA pathway against TEs, LTR retrotransposons exhibit very low expression levels in wildtype *Drosophila melanogaster* ovaries (on average below 0.8 transcripts per million [TPM]; Fig. 1B). Notably, *copia* stands out as an exception, displaying considerable levels of poly-adenylated transcripts exceeding 60 TPM (Fig. 1B), 10x higher than the second highest expressed element *412* (Fig. 1B). To explore the cell-type specificity of this observation, we used a single-cell RNA sequencing (scRNA-seq) dataset from *Drosophila* ovaries [28, 29] and re-analyzed it using a transposon-centric analysis pipeline [30] filtering TEs at a adjusted p-value threshold <= 0.05. This analysis showed that *copia* transcripts were higher in somatic cells of the ovary compared to the germline (Fig. 1C, D). Unlike other LTR retrotransposons that often exhibit cell type-specific expression in the ovarian soma or germline [31], *copia* was strongly expressed throughout the ovarian somatic compartment, including late-stage follicle cells (Fig. S1).

To determine the precise spatio-temporal expression of *copia* in the intact ovary, we performed single-molecule fluorescence *in situ* hybridization (smFISH) experiments. *copia* transcripts were readily detectable in somatic follicle cell nuclei and cytoplasm throughout oogenesis (Fig. 1 E-F, Fig. S2A). In contrast, they were virtually absent in germline cells, where we only detected discrete foci in nurse cell nuclei, likely representing nascent *copia* piRNA precursor transcripts (Fig. S2A). Consistent with the smFISH analysis, immunofluorescence experiments using a polyclonal antibody against *copia* Gag (Fig. S2B) showed signal specifically in the follicular epithelium (Fig. 1G). In addition to a diffuse cytoplasmic signal, we detected immunofluorescence foci within follicle cell nuclei and their periphery (Fig. 1H). Expression of three different shRNA constructs targeting the *copia* genomic RNA within the *gag* coding sequence in follicle cells resulted in reduced cytoplasmic RNA FISH and Gag immunofluorescence signal (Fig. S2C). This observation supports the specificity of our *copia* detection assays.

Many LTR retrotransposons form virus-like particles (VLPs) [32–35]. VLPs have also been described for *copia* in the nucleus of cultured, embryonic macrophage-like *D. melanogaster* S2 cells [36]. In *copia*, VLPs form by encapsidation of the full length gRNA within a capsid formed by proteins expressed from full-length (Fig. 1I, Isoform 1) and spliced (Fig. 1I, Isoform 2) *copia* RNA [37]. To explore whether *copia* forms VLPs in ovarian follicle cells, we sectioned high-pressure frozen, freeze-substituted and resin-embedded stage 10 egg chambers at 300 nm thickness and collected electron tomography data (Fig. 1J-L, Fig. S3). We observed abundant, clearly distinguishable spheres indicative of VLPs (Fig. 1K, Fig. S3B-F), ∼50 nm in size, resembling the previously described *copia* VLPs [36]. The VLPs were rare in the cytoplasm but abundant in the nucleus where they formed large clusters of capsids that were excluded from nucleoli (Fig. 1K, Fig. S3C, E-F). Of note, we detected instances of *copia* VLPs in the vicinity of nuclear pores (Fig. 1L inset). Taken together, *copia* is expressed in ovarian follicle cells and forms abundant, predominantly nuclear VLPs. Consequently, the follicle cell epithelium represents a unique and promising opportunity to study the replication cycle of an abundant and active LTR retrotransposon in its natural host environment.

### The in-cell structure of *copia* VLPs

To visualize *copia* VLPs in intact host cells, we turned to on-grid cryo-correlative light and electron microscopy (cryo-CLEM) experiments using follicle cells isolated from egg chambers. Isolated follicle cells can be vitrified by plunge freezing due to their small cell size (<10 µm) (Fig. 2). To identify the respective cells, we fluorescently labeled somatic ovarian cells using a UAS-controlled Sec13::EosFP transgene and the *traffic jam* (*tj*)-Gal4 driver line, which is active in all somatic cells of the ovary. Follicle cells isolated by dissociation and filtration (Fig. S4, Materials & Methods) showed similar localization patterns of Sec13::EosFP compared to cells in intact egg chambers (Fig. S4 B-C). Cells positive for Sec13::EosFP were identified on frozen-hydrated grids using cryo-fluorescence light microscopy (cryo-FLM) data that was subsequently correlated with the images collected in the FIB-SEM instrument (Fig. 2 A-B, Fig. S4D-J). SEM imaging of the lamella surface was used to target the nucleus for final lamella preparation (Fig. S4H). Low magnification overviews of these lamellae revealed clearly visible nuclear VLP clusters (Fig. 2C). As already seen in the plastic-embedded sections (Fig. 1 J-L), the VLPs were excluded from the nucleolus, identified by the missing Sec13::EosFP signal (Fig. S4J). Cryo-ET tomograms of *copia* VLPs in nuclear clusters (Fig. 2D) showed typical features of retrotransposons [33] such as mature (Fig. 2E) and immature (Fig. 2F) capsid structures. As determined by visual inspection, most of the VLPs (>99%) showed an immature capsid structure.

**Figure 2:**
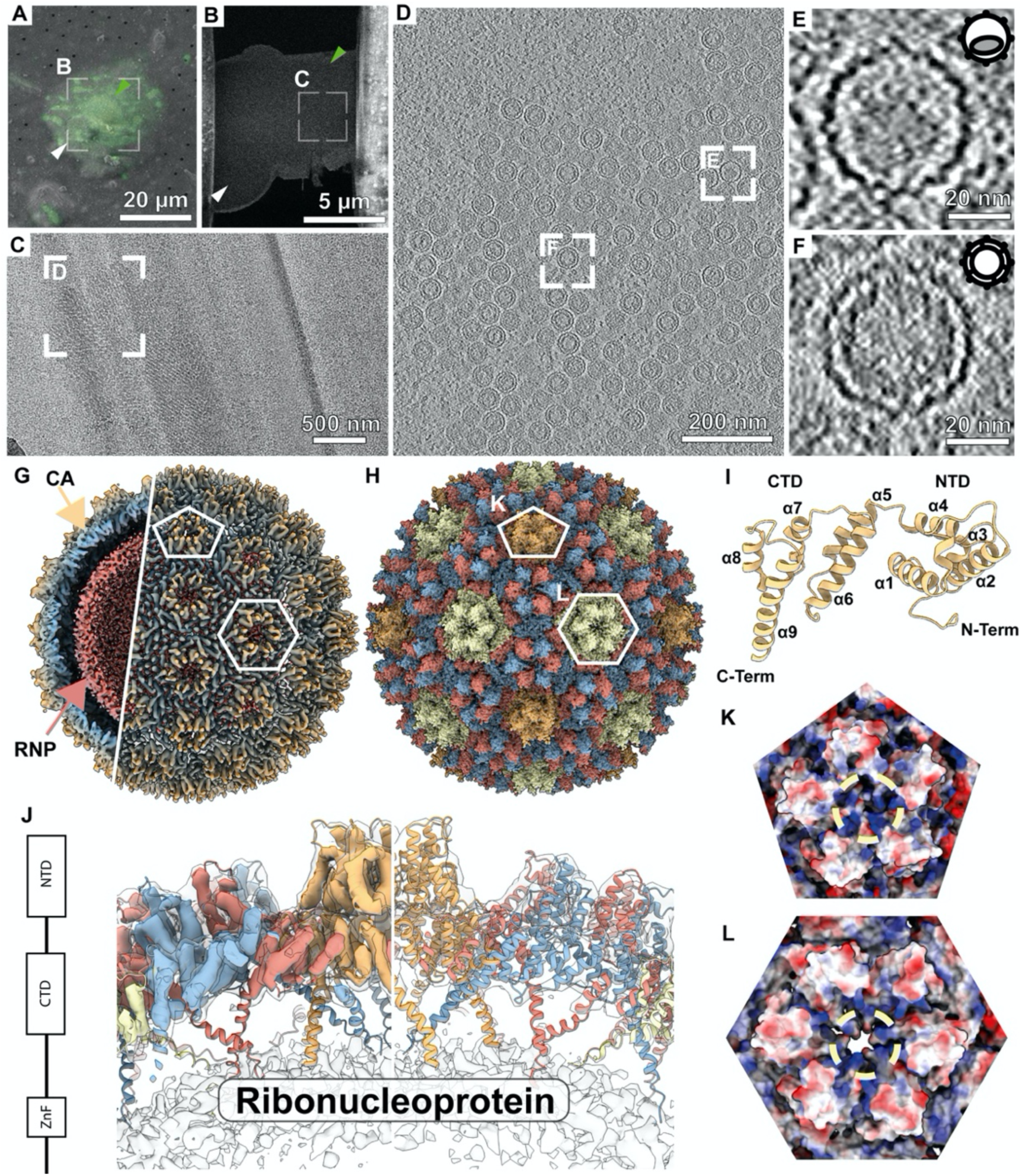
The in-cell structure of *copia* VLPs. **A:** Cryo-SEM image with overlaid fluorescence signal of a two-cell cluster, one negative for Sec13::EosFP (white arrow) and one positive for Sec13::EosFP (green arrow) signal. **B:** SEM image of final lamella prepared from the position in A with arrowheads indicated the Sec13::EosFP positive (green) and negative (white) cells. **C:** TEM overview image showing nuclear cluster of *copia*. **D:** Tomographic slice of nuclear *copia* cluster. Rectangles show zooms into particles shown in E and F. **E:** Tomographic slice of a mature capsid. **F:** Tomographic slice of an immature capsid. Cartoons show a simplified schematic representation of a mature (E) and an immature (F) capsid **G:** Subtomogram average of the entire immature *copia* capsid at 7.7 Å resolution. Note that the capsid and the luminal view of the VLP are displayed at different thresholds. The ribonucleoprotein density only appears at a higher threshold compared to the capsid structure. The structure is colored radially from red (low radius) to blue (medium radius) to yellow (high radius). **H:** AlphaFold-Multimer and molecular dynamics flexible fitting-based model of the entire capsid structure. Depicted are CA conformations of the C5 (orange), C3 (beige), and C1 environments (red and blue). **I:** Structural model of the *copia* CA NTD and CTD. **J:** Side view of a 5-fold symmetric subregion indicated by a pentagon in G and H. Left side shows the density at high threshold colored based on the corresponding subunit, right side shows the model within the low threshold structure. **K:** Top view of the pentameric region shown in B as a surface representation color coded by charge from negative (red) over neutral (white) to positive (blue). **L:** Top view of the pseudo-3-fold symmetric region (hexamer) shown in B as a surface representation color coded by charge as in K. Genotype for all images: *tj > EosFP::Sec13*.

We conducted subtomogram averaging on individual VLPs from nuclear clusters to elucidate the structure of the *copia* capsid. Capsids were identified through template matching, utilizing a hollow sphere as a reference (Fig. S5). In an initial C1 alignment of *copia* capsids, pentameric and hexameric structures emerged, suggesting icosahedral symmetry of the *copia* immature capsid with a triangulation number of T=9 (Fig. S6). Through refinement and classification in Relion and M, approximately 40% of capsids were retained in the final class that yielded the capsid structure at 7.7 Å resolution (GSFSC) (Fig. 2G, Fig. S7, Movie S1). At this resolution, the cryo-EM map displayed discernible alpha-helical features. We used the AlphaFold-Multimer predicted penta- and hexameric structures of the first 267 residues of the *D. melanogaster copia* Gag protein (Fig. S8) as a starting point for molecular modeling (Fig. 2I). The resulting models displayed rotational symmetry formed by 5 (Fig. S8A) or 6 monomers (Fig. S8D). A simple rigid body fit of the predicted pentameric (Fig. S8B-C) and hexameric (Fig. S8E-F) assembly yielded good starting conformations for modeling and allowed for the construction of a full capsid model through manual refinement by interactive molecular dynamics flexible fitting (MDFF) (Fig. 2H-J). Surface models with electrostatic mapping highlighted a highly positively charged central region in both pentameric (Fig. 2K) and hexameric (Fig. 2L) interfaces, seemingly forming a pore in the hexamer that is absent in the pentamer (Fig. S9A-D). While not resolved in the cryo-EM map, the regions of fitted structures constituting the anticipated nucleic acid-binding zinc finger motifs (Fig. S9E-F) pointed towards the putative RNA and NC density on the luminal side of the capsid (Fig. 2J).

The abundance of *copia* VLP clusters (Fig. 1J-L, Fig. 2D, Movie S2) in the nucleus raises the question whether specific protein-protein interactions underlie their formation. Subtomogram averaging allows for placing obtained structures into the cellular context and consequently enables the determination of the spatial relationships between macromolecular complexes. We therefore explored the structural details underlying nuclear cluster formation of *copia* VLPs (Fig. 3A) and identified a repeating structure (Fig. 3B). The clusters deviate from a hexagonal close packing expected for tightest packing of spheres. A neighbor map analysis from subtomogram averaging shows that within the clusters, each particle is surrounded by 12 neighboring particles (Fig. 3B) in what resembles a rhombohedral Bravais lattice with similar angles alpha≈beta≈gamma = 61.5 ° and distances a≈b≈c = 52 nm between all particles (see Materials & Methods). The *copia* assemblies thus resemble a paracrystalline lattice with high mosaicity (Fig. S10A). Consequently, to address whether specific inter-capsid interactions underlie the formation of *copia* clusters, we performed a nearest neighbor analysis for the C1 (also called pseudo-C3), C3, and C5 environments of the icosahedral lattice (Fig. 3C). Focusing on events with a cutoff distance of 3.6 nm based on the smallest distance distribution (Fig. S10B) yielded a distribution of contacts that differed from a distribution for randomly oriented VLPs (Fig. 3D). Especially C1-C5 and C5-C5 contacts with measured distribution of 56 % and 13%, respectively, were strongly overrepresented compared to a distribution of 18% ± 2 % and 2% ± 1 %, respectively, for randomly oriented VLPs (Fig. 3D). Subtomogram averaging of the contact positions yielded reconstructions suggesting a preferred inter-capsid distance and orientation (Fig. 3E-J, Fig. S10C). The average is especially clear for the C5-C5 (Fig. 3E-F) and C1-C5 (Fig. 3I-J) contacts, which also show the highest deviation in occurrence from randomly oriented VLPs (Fig. 3D). Mapping the three predominant contacts C1-C5, C5-C5, and C1-C1 in the lattice (Fig. 3D) between VLP positions illustrates the interactions present in the nuclear *copia* clusters (Fig. 3K, Fig. S10D-L, Movie S3).

**Figure 3:**
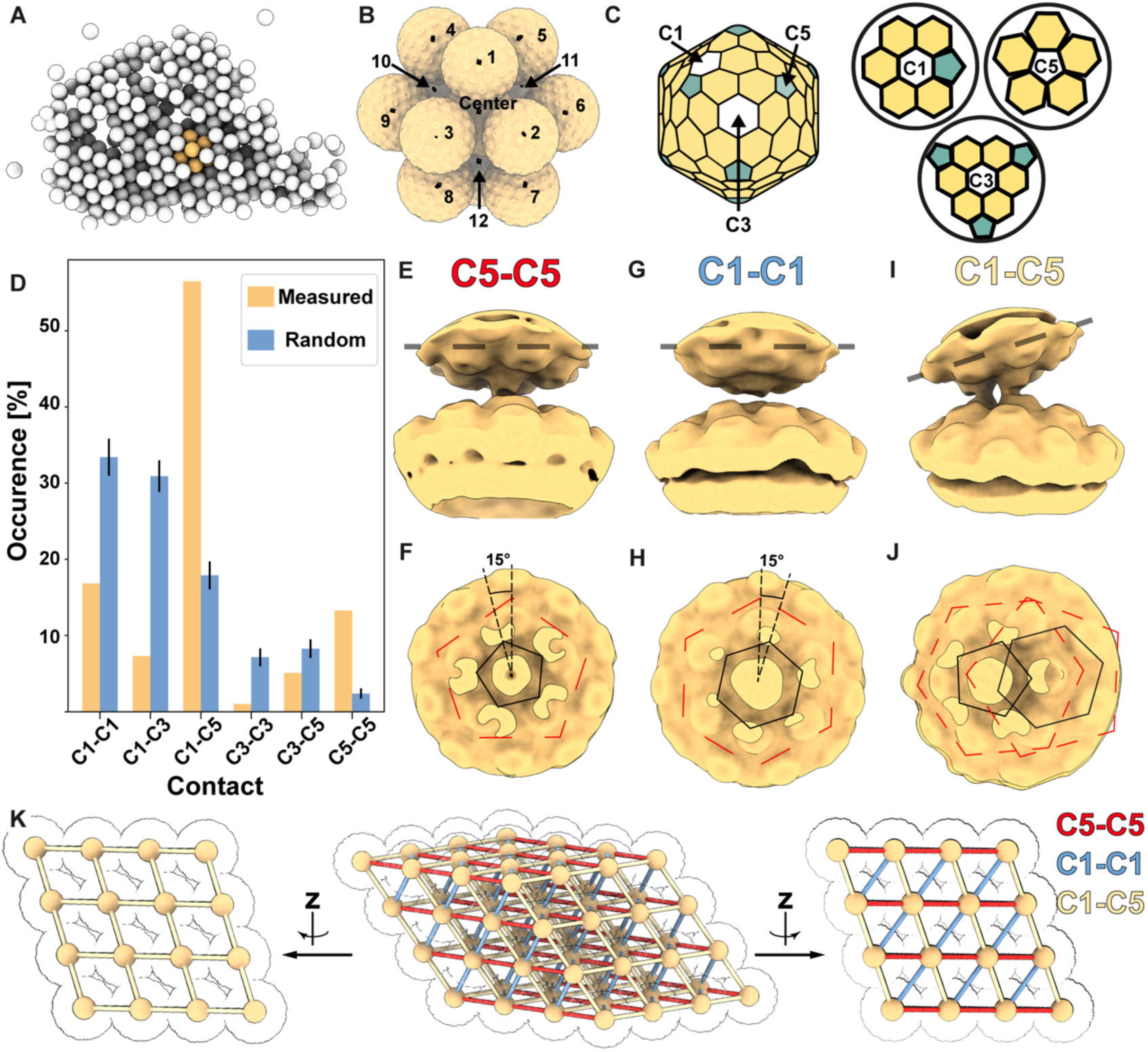
Cluster formation of *copia* VLPs in the nucleus. **A:** Rendering of a nuclear *copia* cluster. Highlighted in yellow is one of the repetitive units of capsid positions central capsid with 12 neighbors as determined by the template matching scoring map. **B:** Zoom on a repetitive unit from the particle positions in A overlaid with the template matching neighbor volume highlighting the 12 neighbors relative to the central capsid. **C:** Schematic representation of the different environments C1 (also called pseudo-C3), C3, and C5 present in an icosahedral lattice with triangulation number T=9. **D:** Comparison of contact distribution between the measured contacts to a random orientation of icosahedral particles. **E:** Side view of the reconstruction of C5-C5 contacts. Dashed line indicates the cross-section plane shown in F. **F:** Cross-section through the C5-C5 contact. **G:** Side view of the reconstruction of C1-C1 contacts. Dashed line indicates the cross-section plane shown in H. **H:** Cross-section through the C1-C1 contact. **I:** Side view of the reconstruction of C1-C5 contacts. Dashed line indicates the cross-section plane shown in J. **J:** Cross-section through the C1-C5 contact. Red polygons are the environments within the image plane, while black polygons are the environments above the image plane (see also Fig. S10G-L). **K:** Model of extended lattice based on experimental positions shown in three orientations with contacts represented as sticks colored according to C5-C5 (red), C1-C1 (blue), and C1-C5 (yellow) contacts. Additionally, the particle centers (yellow spheres) and corresponding capsids (black outline) on the lattice are shown.

### Nuclear VLPs are small and more homogenous

To visualize *copia* VLPs in intact *D. melanogaster* egg chambers (Fig. 4A), we performed cryo-lift-out [25, 38] experiments on high-pressure frozen egg chambers (Fig. 4B, Fig. S11), with a particular focus on the nuclear periphery of follicle cells.

**Figure 4:**
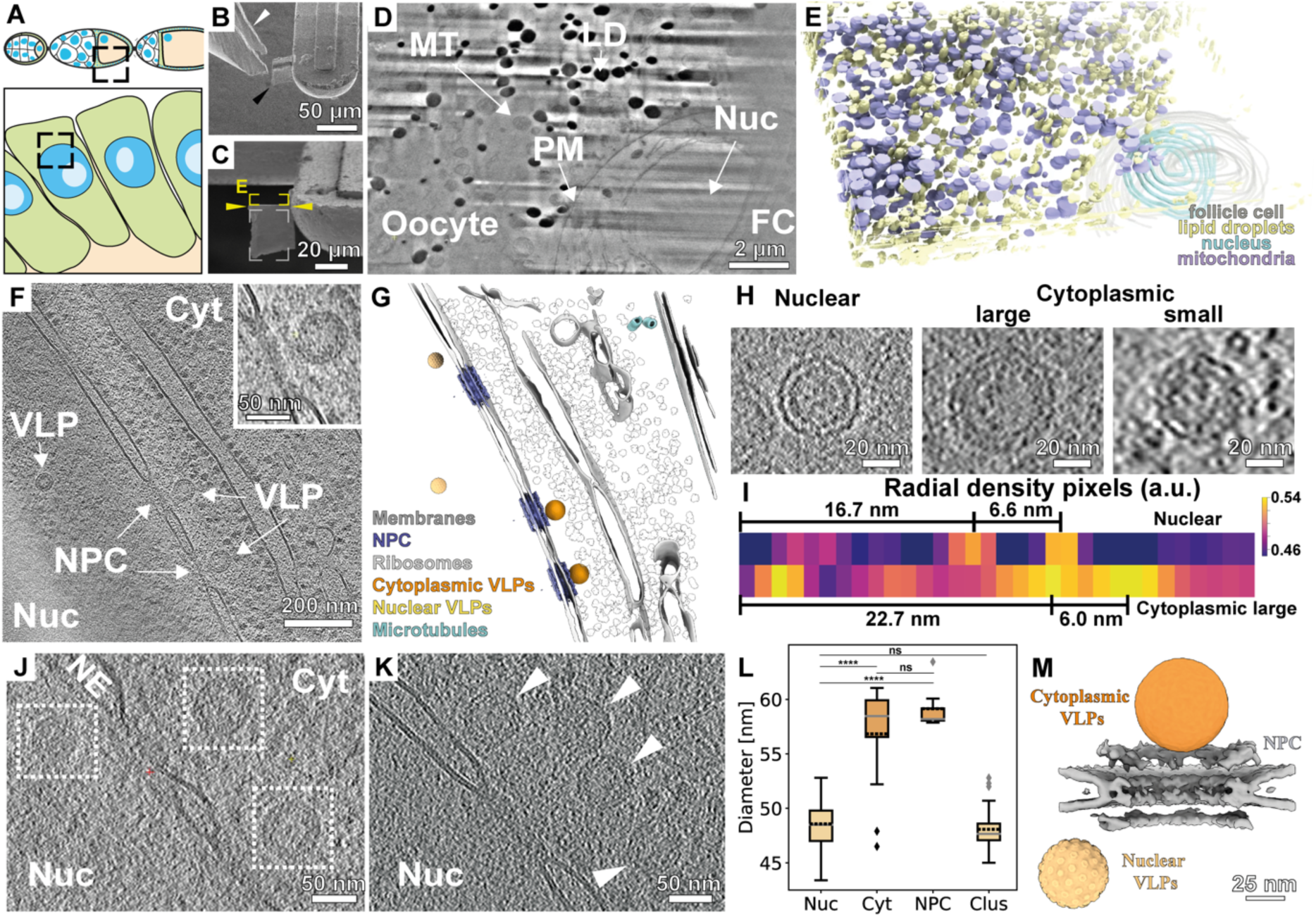
Cryo-lift-out on intact *D. melanogaster* egg chambers. **A:** Schematic representation of the area targeted by cryo-FIB lift-out. **B:** SEM image of the large block of material (black arrowhead) transferred to a receiver grid by cryo-lift-out using a piezo-driven tweezer (white arrowhead). **C:** Focused ion beam image of the lift-out volume. Yellow rectangle indicates the volume investigated during FIB-SEM volume imaging for targeting the final lamella, yellow arrowheads show the chosen position for the final lamella and the white rectangle the volume removed from below during lamella preparation. **D:** SEM image of lift-out volume showing mitochondria (MT), lipid droplets (LD), the plasma membrane (PM) of a follicle cell (FC) and its nucleus (Nuc). **E:** 3D rendering of the volume obtained by FIB-SEM volume imaging. **F:** Tomographic slice from a lift-out experiment showing virus-like particles (VLPs) engaged with the cytoplasmic (Cyt) side of nuclear pore complexes (NPC) and a VLP in the nucleus (Nuc). Inset shows a zoom on a nuclear pore complex with a VLP aligned to its central channel. **G:** 3D segmentation of **F**. **H**: To-scale comparison of VLPs found in the nucleus and cytoplasm. A large and small example is shown for the cytoplasmic VLPs. **I**: Normalized radial density plots of 5 particles each shown for nuclear and large cytoplasmic VLPs. **J**: Tomographic slice showing two cytoplasmic VLPs aligning with the NPC’s central channel and a nuclear VLP in NPC proximity on the nuclear side. **K:** Tomographic slice of a cluster of VLPs at an NPC. **L:** Quantification of VLP sizes classified into 4 regions, single VLPs in the nucleus (Nuc, n = 37), in the cytoplasm (Cyt, n = 14), aligned to the central channel (NPC, n = 7), and VLPs in nuclear clusters (Clus, n = 40). Dashed black lines show the mean, solid grey lines the median of the distribution. Unpaired Student’s t-test was performed to test significance. Ns: p > 0.05. *: p ≤ 0.05, **: p ≤ 0.01, ***: p ≤ 0.001, ****: p ≤ 0.0001. **M:** 3D rendering of the approximate VLP positions shown in J for size comparison. Genotype of all images except “cytoplasmic small” in I: *LifeAct::GFP*. Cytoplasmic small: *tj>Sec13::EosFP*.

We targeted follicle cell nuclei by using cryo-FIB-SEM volume imaging of large lift-out volumes from the top surface of the attached biological lift-out material (Fig. 4C). Once the nucleus became discernible in the SEM micrograph (Fig. 4D), the final lamella was prepared from below. This methodology not only ensured that the lamella contained the nucleus, but also provided contextual insight into the volume around the prepared thin section, which was amenable to 3D visualization through segmentation (Fig. 4E). The acquired tomograms of the nuclear periphery within follicle cells revealed the presence of both nuclear and cytoplasmic VLPs (Fig. 4F-G, Movie S4). We found that nuclear capsids free floating (48.6 nm ± 1.8 nm in diameter, n = 37) and in clusters (48.1 nm ± 1.7 nm, n = 40) displayed a similar capsid size in both intact egg chambers and isolated cells (Fig. 4H). Cytoplasmic VLPs showed predominantly large capsid structures with small outliers (56.8 nm ± 4.5 nm, n = 14) (Fig. 4H, L). Similar results were observed using radial average density profiles (Fig. 4I, Fig. S12). Several VLPs were situated in such close proximity to the NPC that they are in reach of central channel nucleoporins [39] and dip into the typical ribosome exclusion zone observed around nuclear pores [40] (Fig. 4F inset, Fig. 4J-M). The diameters of VLPs in reach of central channel nucleoporins was slightly larger (59.1 nm ± 1.89 nm, n = 7) (Fig. 4L) and no small capsids were observed at the cytoplasmic side of the NPC. The smaller capsid diameters are consistent with the T=9 *copia* capsid structure determined above (Fig. 2, Fig. S12). Larger assemblies would instead be consistent with a higher triangulation number of the capsid, namely T=13 capsids encompassing 780 proteins, which would exhibit a theoretically size of 13^½^ * 9^-½^ * 48.6 nm = 58.4 nm based on the T=9 capsid diameter from radial density plots.

### *Copia* is differentially repressed in somatic cells and the germline

In metazoans, invading TEs need to reach the host’s germline for vertical transmission to the progeny [41] and successful transposons often show germline-restricted expression [42]. Therefore, the abundant expression of *copia* in somatic cells was surprising. Unlike Errantiviruses, a distinct subgroup of LTR retrotransposons with a known capacity for soma-to-germline transmission via enveloped VLPs [31, 43, 44], *copia* lacks an *envelope* (*env*) open reading frame. Sensitive single-molecule RNA FISH experiments confirmed the absence of *copia* sense transcripts in the growing oocyte of wildtype ovaries (Fig. S2A), arguing against a non-canonical, *env*-independent soma-to-germline transmission process. These data are consistent with the absence of an anti-*copia* defense program by the host piRNA pathway in follicle cells that efficiently silences the expression of Errantiviruses [45–47]: piRNAs targeting *copia* in somatic ovarian cells are approximately 10- to 100-fold lower than those targeting the different Errantiviruses (Fig. 5A). In alignment with this, *copia* expression was only moderately increased, ∼2-4-fold in ovaries with a disrupted somatic piRNA pathway [31], also discernible in smFISH experiments (Fig. 5B).

**Figure 5:**
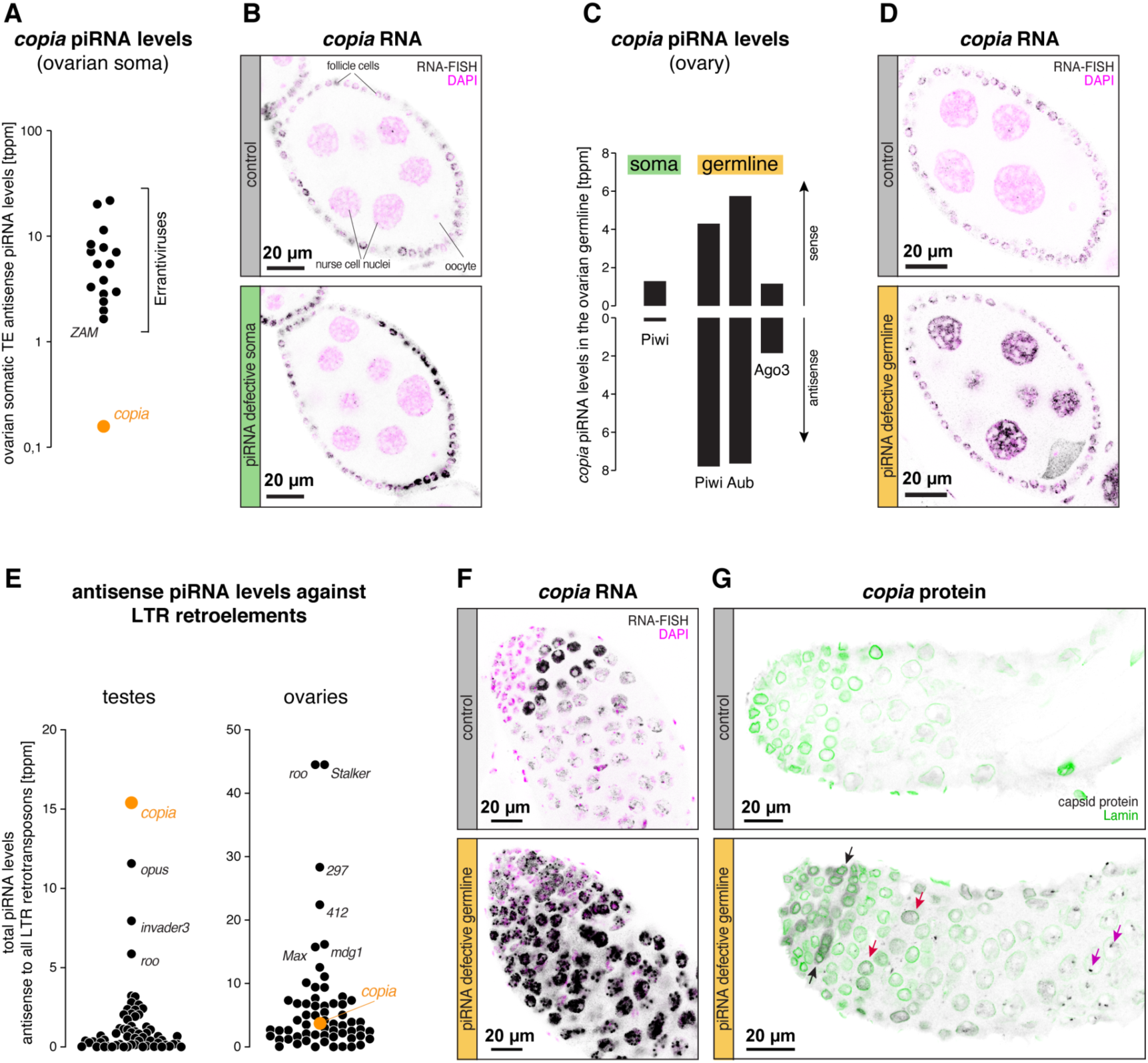
*Copia* regulation and localization in ovaries and testes. **A:** Jitter plot comparing the levels of soma Piwi bound antisense piRNAs mapping with zero mismatches to Errantiviruses (black) that are strongly repressed in somatic follicle cells and *copia* (orange). **B:** Confocal images show *copia* smFISH (black) and DAPI (magenta) in stage 6/7 egg chambers of soma specific control and piRNA pathway knockdowns. Control: *tj>arr2/GD*; piRNA defective soma: *tj>vreteno/GD*. **C:** Plot summarizing levels of piRNAs bound to somatic Piwi, and bound to germline Piwi, Aub and Ago3 mapping to *copia* in the ovary with zero mismatches. **D:** Confocal micrographs showing *copia* smFISH (black) and DAPI (magenta) stage 6/7 egg chambers of germline specific control and piRNA pathway knockdowns. Control: *MTD>white sh*; piRNA defective germline: *MTD>aub ago3 sh*. **E:** Jitter plots displaying the levels of anti-sense piRNAs from total small-RNA-seq of *white*^1118^ ovaries and testes mapping with zero mismatches to *copia* (orange) and all other *Drosophila* LTR retrotransposons (black). **F-G:** Confocal images showing germline specific control and piRNA pathway knockdowns in testes. Control: *NGT40&bam>white sh;* piRNA defective germline: *NGT40&bam>aub ago3 double sh*. **F:** *Copia* smFISH in black and DAPI in magenta. **G:** Localization of *copia* Gag (black) and LaminDM0 (nuclear outline, green). Black arrows indicate cytoplasmic *copia* protein localization in spermatogonia, red and magenta arrows highlight dispersed and concentrated nuclear *copia* signal in early and differentiating spermatocytes, respectively. Control: *NGT40; bam-GAL4>white sh*; piRNA defective germline: *NGT; bam-GAL4>aub ago3 sh*.

In contrast to the ovarian soma, *copia* antisense piRNAs are abundant in the germline and display the characteristic features of a canonical germline piRNA program (Fig. 5C, Fig. S14). These features include (1) a robust ping-pong signature evident through sense/antisense piRNA pairs bound to the germline-expressed PIWI-clade Argonaute proteins Aubergine (Aub) and Ago3 (Fig. S14A), and (2) a diverse repertoire of Piwi-bound piRNAs (Fig. S14B). To assess the impact of the piRNA pathway on *copia* expression in the female germline, we depleted ovaries of Aub and Ago3 in the germline using transgenic RNA interference (Fig. S14C). This resulted in a ∼30-fold decrease in *copia* antisense piRNA levels [48]. Concomitantly, polyA-plus RNAseq experiments revealed an 11-fold increase in ovarian *copia* sense transcripts. *copia* transcripts exhibited a moderate accumulation in the growing oocyte but were highly enriched in nurse cell nuclei (Fig. 5D, Fig. S14C). This differed from other germline-adapted LTR retrotransposons in ovaries whose transcripts accumulated strongly in the oocyte of piRNA defective ovaries (e.g. *McClintock,* Fig. S14D). Notably, despite increased *copia* sense transcript levels, and in contrast to the *McClintock* retrotransposon, we did not observe increased levels of *copia* Gag protein in the germline of piRNA-deficient ovaries (Fig. S14C). *Copia* thus shows limited adaptation for propagation within the ovarian germline, setting it apart from other LTR retrotransposons adapted for the female germline [31, 48].

Prior investigations of *copia* have revealed a predominant amplification of genomic *copia* copies within the male germline [49, 50]. Notably, *copia* antisense piRNAs are the predominant species in testes when all 62 LTR retrotransposon lineages are considered (Fig. 5E, Fig. S15A). In comparison, the piRNA repertoire in the female germline shows moderate antisense piRNA levels against *copia* compared to other LTR retrotransposons (Fig. 5E, Fig. S15B). This observation argues that the *Drosophila* host has evolved a particularly stringent piRNA control mechanism against *copia* in the male germline, supporting a replication cycle of *copia* within the testes niche. Experimental depletion of Aub and Ago3, the two dominant PIWI-clade proteins responsible for TE silencing in testes [51], led to a strong increase in *copia* sense transcripts, as evidenced by RNA FISH experiments: In wildtype testes, *copia* sense transcripts were detected only within the nuclei of spermatocytes, with negligible presence in spermatogonia and differentiating spermatocytes (Fig. 5F). Conversely, depletion of the piRNA defense system in the male germline resulted in a pronounced increase of *copia* FISH signal throughout spermatogenesis (Fig. 5F). *copia* transcripts were present in the cytoplasm of spermatogonia and accumulated within the nucleus, often in discrete foci, of maturing spermatocytes. Immunostaining with the *copia* Gag antibody corroborated these findings, revealing abundant *copia* protein only in piRNA-defective testes: *copia* protein initially accumulated in the cytoplasm of spermatogonia and concentrated in the nuclei of spermatocytes during later stages (Fig. 5G).

Our comprehensive *in vivo* analysis suggests that *copia* has evolved several traits for an amplification in the male germline: a promoter with strong activity in germline cells as apparent from the Aub-Ago3 knockdowns (Fig. 5D, F), male-specific translation of the Gag protein (Fig. 5G, Fig. S14C), and translocation of the Gag protein (Fig. 5G), potentially as VLPs, into the nucleus. The latter trait is particularly notable as efficient nuclear translocation of *copia* VLPs would be counterproductive in the developing female germline: Here, LTR retrotransposon capsids avoid being imported into nurse cell nuclei from where they are expressed but instead hijack the microtubule network connecting nurse cells and oocyte in order to target the genome of the growing oocyte [31, 52].

## Discussion

In this study, we provide a comprehensive investigation of the in-cell structure and snapshots of the replication cycle of the *copia* retrotransposon in *D. melanogaster*. We elucidate the *copia* capsid structure at subnanometer resolution inside ovarian somatic cells and report the subcellular localization of *copia* VLPs, using a variety of imaging techniques. The *copia* Gag protein forms an icosahedral capsid structure (T = 9) comprising 540 monomers in a conformation resembling the mature CA structure of HIV. It confirms the previous observation for Ty3 that immature retrotransposon capsids resemble mature retrovirus structures [33]. We further observe VLPs *in vivo* with a size distribution from ∼48 nm to ∼58 nm that are consistent with T = 9 and T = 13 *copia* capsid assemblies. The structural model for the T = 9 *copia* capsid density shows pores within the hexameric assemblies that are positively charged, which are filled in the pentameric assemblies (Fig. 2K-L, Fig. S9A-D). Similar pores have been described to be relevant for nucleotide import during reverse transcription of HIV-1 [53]. On the luminal side of the capsid, an additional density likely represents the *copia* ribonucleoprotein. In the AlphaFold prediction, helices extend into that density, suggesting the presence of a nucleic acid binding site close to the C-terminus of *copia* CA. Indeed, previous description of the *copia* sequence [7] place a CCHC-type zinc finger motif at residue 230-247 of the full length *copia* Gag-IN-Pol (GIP) polypeptide (Fig. S9E-F). While not resolved in the subtomogram averaging density, those residues would coincide with the luminal density of the immature capsid structure based on our structural model (Fig. 2J).

Clustering of retrotransposon capsids has previously been observed for *copia* in *D. melanogaster* S2 cells [36] as well as for other retroelements [33, 35, 54, 55]. However, the structural or molecular basis of this phenomenon is unknown. Using cellular cryo-ET we were able to describe the interparticle relationship of the *copia* capsid paracrystalline arrays. We find that cluster formation is driven by inter-capsid interactions that differ from randomly oriented closed hexagonal packing of VLPs (Fig. 3). Whether cluster formation is relevant to the replication cycle or regulation of *copia* integration remains to be determined. We show, however, that there are preferred capsid-capsid interactions that promote clustering. In conjunction with the presence of clustering in various retrotransposons, clustering is likely carrying an evolutionary advantage.

We found only 1.1 % of the *copia* VLPs to localize to the cytoplasm (21 out of 1979 from ∼50 tomograms that show VLPs). The low number of cytoplasmic particles observed, which is similar to previous estimates of 0.4% [56], raises the question of what cellular mechanisms drive this distribution. For other retrotransposons, including the *copia*-related element Ty1 in *S. cerevisiae* [57], capsid assembly in the cytoplasm has been shown [35, 58]. Given the low number of detectable cytoplasmic particles despite the abundance of *copia* RNA in the cytoplasm [59], which is required for capsid assembly, we hypothesize that cytoplasmic assembly followed by efficient transport into the nucleus occurs (Fig. 6A). While we formally cannot rule out that capsid assembly could also occur within the nucleus, several findings would be in line with a model in which *copia* VLPs can enter the nucleus intact and that nuclear import may be a VLP size selector and, thus, facilitator of nuclear cluster formation:

- Structural relationship to HIV (HIV CA hexamer binds FG-Nups)
- Size of the smaller capsids compatible with nuclear import
- Enrichment in the nucleus

**Figure 6:**
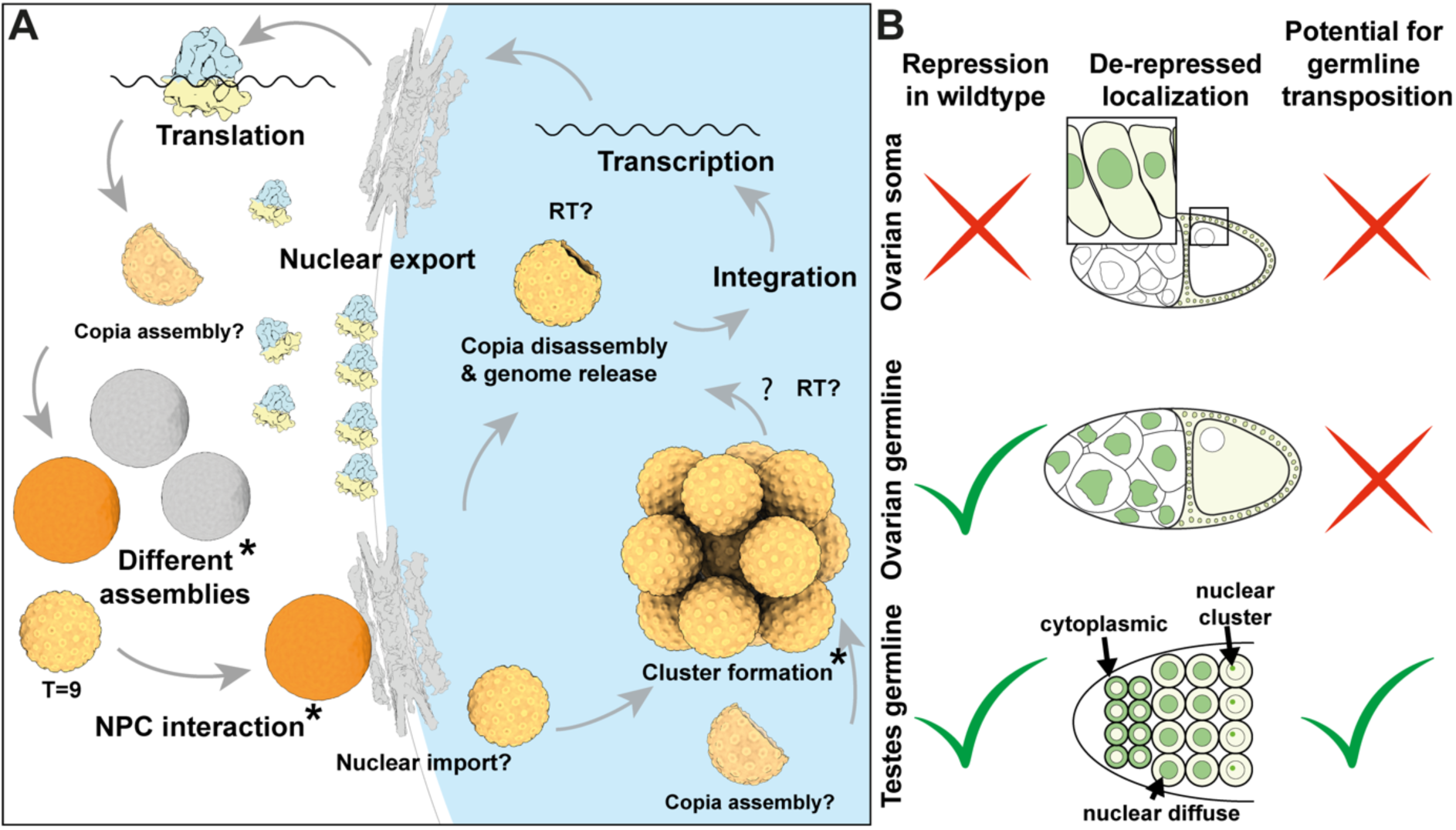
Proposed model of the *copia* replication cycle and its transposition. **A:** Model of the *copia* replication cycle. Asterisks indicate states that were observed by cryo-ET. **B:** Summary of the repression, localization (green representing detected signal for *copia* in the different compartments), and proposed potential for germline transposition of *copia* expression in ovarian soma, ovarian germline, and testes germline.

Another finding in support of this hypothesis is that we observe large *copia* capsids that align with the central channel of the NPC, suggesting an interaction with the nuclear pore. Such an NPC interactions would be especially interesting in light of recent findings of HIV capsids being transported as a whole through the NPC [14] by mimicking karyopherins [15, 16]. The observations presented here further support that other retroelements and retroviruses may undergo NPC-mediated nuclear translocation in fully assembled capsids. It should be noted, however, that while the CA fold and mature CA structure is highly conserved among retroviruses and retrotransposons, NPC-mediated translocation will only manifest in a subset of these elements as some retroviruses (e.g. MuLV) have been shown to require passage through mitosis for nuclear entry [60].

It is surprising to find *copia* VLPs in such high numbers in the somatic nuclei of the egg chamber (Fig. S1) [36] as their rampant integration would pose a threat to genome integrity. It is possible that additional factors are required for successful integration of new *copia* copies. Somatic expression of transposition incompetent *copia* might also be selected for due to benefits for the host. It has recently been proposed that *copia* could act as regulator of plasticity at synapses and more specifically at the neuromuscular junction (NMJ) of developing *Drosophila* larva, as RNAi-mediated knockdown of *copia* resulted in an increase in the formation of boutons, synaptic swellings at the end of axons [61].

Irrespective of a potential biological function in the soma, *copia* transposition in the immortal germline genome is essential for its evolutionary survival. *copia* transposition has been previously shown to be limited to the male germline [50]. Due to this germline-specificity, an alternative piRNA suppression mechanism involving spermatogenesis-specific events was proposed [62]. While remaining speculative, our *copia* VLP cryo-ET data, coupled with the observed expression of first cytoplasmic and then nuclear *copia* Gag protein in piRNA pathway deficient testes (Fig. 5G), suggests that the event restricting *copia* transposition to the male germline may indeed be the nuclear entry of *copia* VLPs into spermatogonia (Fig. 6B). We have shown here that the integration of cryo-ET inside cells and tissues with multimodal data can play a vital role in answering some of the remaining questions around the replication cycle of LTR retrotransposons. Our study demonstrates how the combination of imaging modalities and high-resolution imaging by cryo-ET helps to assess the ‘molecular sociology’ [63] of LTR retrotransposons, and *copia* specifically, in their relevant host environment at molecular resolution.

## Acknowledgement

We thank Beata Turonova for fruitful discussion on image processing, Jürg Müller for fly room access and discussions, Ben Engel, Anne Schütz, Philipp Erdmann, Davide Torre, and James Stacey for discussions. We are grateful to Jean-Marc Philippe in the laboratory of Thomas Lecuit who made the Sec13::EosFP construct. We thank Lazlo Tirian and Ralf Jansen for technical support, Deeptiman Chatterjee for help with scRNAseq analysis, and Stefanie Böhm for critical reading of the manuscript. This study used the infrastructure of the Department of Cell and Virus Structure at the MPI of Biochemistry and EMBL EM core facilities (EMCF). S.K. was supported by the International Max Planck Research School for Molecular and Cellular Life Sciences. This work was funded by the Max Planck Society (JMP, JAGB, and MB).

## Code and Data availability

Plastic tomograms of *D. melanogaster* egg chambers are deposited under EMDB accession codes: EMD-19643, EMD-19644, EMD-19645, EMD-19646. Example cryo-ET tomograms from isolated cells (EMD-19647) and lift-out lamellae (EMD-19648) were uploaded to EMDB. The subtomogram averages of the nuclear *copia* capsid structure and a structural model are available under the EMDB and PDB accession codes EMD-19708 and 8S41, respectively. Contact maps (C1-C1: EMD-19649, C1-C5: EMD-19650, C5-C5: EMD-19651) have been deposited. All deposited data will be released upon publication. The code for RNAseq analysis and the Seurat object are available on github: https://github.com/sklumpe/copia-manuscript. Other data is available upon request.

## Author contributions

S.K., K.S., B.H., J.S., performed *Drosophila* experiments, S.K. performed cryo-ET experiments. P.R. performed room temperature TEM experiments. S.K. and F.B. analyzed cryo-ET data. A.Y. performed scRNA-seq analysis. S.K. performed molecular modeling. J.A.G.B. provided guidance and expertise on retrotransposons. S.K., J.B., K.S., M.B., and J.M.P. wrote the manuscript with input from all authors. S.K., J.B., M.B, and J.M.P. supervised the project.

## Materials and Methods

### Fly husbandry

*Drosophila melanogaster* fly stocks were maintained at 18-22 °C on standard cornmeal agar in round-bottom vials. Flies were transferred into a fresh vial supplemented with yeast paste 24 h at 25 °C prior to experiments. All RNAi knockdown crosses were performed at 25 °C.

### Fly strains

The following fly lines were used throughout this study: *w; tj-GAL4/CyO;;* [64]; *w* P{UAS-Sec31.RFP}13.5; snaSco/CyO, P{ftz-lacB}E3* (Bloomington #86533); *emGFP::nup358 PBac{y[+mDint2] = vas-Cas9}VK00027* [40]*; w*; P{w+, sqhp>LifeAct::mNeonGreen};* MKRS/TM6. [65] (Gift from Dr. Cristina Pallares Cartes); *w* P{UAS-Sec13::EosFP} / CyO;* (this study); *w;*; *arrestin2/GD* (VDRC# 20991); *w;; vreteno/GD* (VDRC# 34897); y[1] v[1]; P{y[+t7.7] v[+t1.8]=TRiP.HMS00004}attP2/TM3, Sb[1] (Bloomington #33613); P{w[+mC]=otu-GAL4::VP16.R}1,w[*];P{w[+mC]=GAL4-nanos.NGT}40; P{w[+mC]=GAL4::VP16-nanos.UTR} CG6325[MVD1]; (Bloomington #31777); w;; white sh; [48]; w; aub sh (w22, attP40, 2nd chrs); ago3 sh (v22, attP2 third chrs) [48]; w; NGT40 (2); bam-GAL4 (3) [31] and gift from Jeroen Dobbelaere [66].

### Generation of *copia* short hairpin RNAi transgenes

21 nucleotide short hairpins were designed in http://biodev.cea.fr/DSIR/ against the *copia* Gag open reading frame.

TL67: *copia* single sh was cloned into the W20 sh vector as in [67]. TL67 sh sequence was TTTAAGCACATCTTGCTCGGC.

TL63 and TL68 were cloned using the W20 double sh vector as in [68]. TL63 hairpin sequences were: TAACTTTCTTGCTTGTATCGT and TAATAGTGACATCTCACTCGA. TL68 hairpin sequence was 2x TAATAGTGACATCTCACTCGA. All hairpin vectors were sequence verified and were inserted into attP2.

### Generation of pPW-Sec13::EosFP

pPWG, a pUASp GATEWAY destination vector containing an EGFP tag downstream of a GATEWAY cassette, obtained from Terence Murphy (Carnegie Institute), was modified by replacing EGFP with mEOS fluorescent protein, giving a pPWeos destination vector. The Sec13 (CG6773) ORF was inserted into pDONR221 (Life technologies) using BP GATEWAY cloning, resulting in the p221-Sec13 entry clone. The pUASp-Sec13::EosFP cassette was used for standard *Drosophila* transgenesis and F_2_ transgenes were identified by eye color. The final construct controls expression of the Sec13 protein tagged C-terminally with mEos fluorescent proteins.

### Poly(A) RNAseq

Total RNA was extracted with TRIzol from ovaries of adult flies. After RNAeasy column purification with on-column DNAse I digest, eluted RNA was 2x polyA selected using magnetic oligo-dT beads and further processed as strand-specific RNA-seq libraries [69]. The kit <u>NEBNext Ultra II Directional RNA Library Prep Kit for Illumina</u> was used for this procedure.

### Small RNAseq

Previously sequenced small RNA libraries were analyzed as in [48] and [70].

### Single cell RNAseq analysis

SoloTE was used for scRNA-seq analysis since the standard analysis workflow discards features such as transposable elements due to their repetitive nature. We chose soloTE as it is computationally more efficient compared to other tools and additionally keeps the locus-specific position of the identified TEs [30]. Raw data files were obtained from a public data set (Raw data: SRX7814226, Annotations: GSM4363298) [28] and aligned using the STAR aligner STARsolo [71] against the *D. melanogaster* genome (assembly BDGP6.32 [72]). The resulting expression matrix containing the gene-as well as TE-expression intensities were analyzed using Seurat version 5 [73]. The expression matrix was filtered to remove low-quality cells using the default analysis pipeline. Data was normalized and scaled to prevent highly-expressed genes from dominating the results. Dimensionality reduction was performed using a PCA. A KNN-Graph was constructed based on the Euclidean distances in PCA space to identify the clusters of cells. Marker analysis was carried out using the Seurat FindAllMarkers command keeping only the results with logFC ≥ 0.25 and a p-value ≤ 0.01.

### Single molecule FISH

Single molecule fluorescence *in situ* hybridization probes (smFISH) were designed, prepared and experiments performed as described in [48, 74]. smFISH oligos were designed against full length *copia* Open Reading Frame (ORF) (see Supplementary Table).

### Immunofluorescence of ovaries and testes

Antibody stainings were performed as in [48]. In brief, ovaries and testes were dissected from 2-3 day yeast fed flies and fixed in 4% PFA in PBS with 0.3 % Triton X-100 for 20 minutes at room temperature under constant agitation. Dissected tissues were blocked for at least 1 hour in PBS with 0.5% BSA and 0.3% Triton X-100 at room temperature under agitation. All further incubations (with antibodies and for washes) were performed in the same blocking buffer as above. Primary antibodies were incubated with blocked samples under constant agitation overnight at 4 °C. After four 20-minute washes at room temperature, secondary antibodies were incubated overnight at 4 °C or for at least 2 hours at room temperature followed by a 5-minute wash including DAPI and four 20-minute washes prior to mounting in a hard-setting Prolong diamond mounting medium. Antibodies used were: Cusabio rabbit anti-*copia* Gag (at 1/1000), rabbit anti-McClintock Gag polyclonal Antibody raised against peptide ITEAQTAENFRPQASEQANS [31] (at 1/500), mouse anti-Aub monoclonal Ab 8A3-D7 [48] (at 1/1000), rabbit anti-Ago3 polyclonal Ab [74] (at 1/500), and mouse monoclonal ADL84.12 anti-laminDM0 (from DSHB) (at 1/200). Imaging of smFISH and immunofluorescence-stained preparations was performed using Zeiss LSM780 and LSM980 Airyscan 2 confocal microscopes.

### Resin embedding sample preparation and room temperature tomography

*D. melanogaster* egg chambers were dissected from ovaries in Schneider’s medium and subsequently high-pressure frozen (HPM010, AbraFluid) using Schneider’s medium supplemented with 20% Ficoll (70 kDa) as a cryoprotectant [75]. Following high-pressure freezing, the samples underwent freeze-substitution (EM-AFS2 - Leica Microsystems) with 0.1% Uranyl Acetate (UA) in acetone at −90°C for 48 hours. Then, the temperature was increased to −45 °C at a rate of 3.5 °C per hour, followed by a further incubation for 5 hours. After rinsing in acetone, infiltration with Lowicryl HM20 resin was performed with the temperature rising to −25 °C at a rate of 1.6 °C per hour, followed by 3x 10h in 100% resin at −25 °C. UV light for 48 hours at −25 °C and an additional 9 hours with a gradual temperature increase to 20°C (5°C per hour) polymerized the Lowicryl. Thin sections (300 nm) were cut from the polymerized resin block using a FC7/UC7 ultramicrotome (Leica Microsystems, Vienna, Austria) and placed on carbon-coated mesh grids. Subsequent to post-staining, tilt series of the designated areas were collected from −60° to +60° with an increment of 1° at a nominal magnification of 20,000x resulting in a pixel size at the sample of 12.2 Å at bin2 using a TECNAI F30 TEM at 300 kV (FEI, Thermo Fisher Scientific, Eindhoven, The Netherlands) equipped with a Gatan OneView camera. Tomograms were reconstructed and visualized using the IMOD software package [76].

### Follicle cell isolation and plunge freezing

The UAS-Gal4 system [77] was used for tissue specific expression of a fluorescently labelled protein. To generate females expressing Sec13::EosFP in follicle cells, *w* P{UAS-Sec13::EosFP}/ CyO*; flies were crossed to *;tj-gal4/CyO;;* flies, a follicle cell specific Gal-4 driver lines. Adult female F_1_ progeny carrying both transgenes were scored for subsequent experiments.

Follicle cell isolation was performed similar to protocols for single cell sorting [78]. Ovaries were dissected in Schneider’s medium supplemented with 10% fetal bovine serum, 50 unit/ml penicillin, 50 mg/ml streptomycin, 0.25 mg/ml fungizone and 0.4 µM insulin. 10-15 ovaries were dissected into fresh 2 ml Eppendorf tubes and subsequently washed three times with phosphate buffered saline (PBS) containing 0.137 M NaCl, 27 mM KCl, 10 mM Na_2_HPO_4_, 1.8 mM KH_2_PO_4_. The digestion of the tissue was done in 0.5% Trypsin in PBS. First, the oocytes were sedimented and the supernatant was removed. Subsequently, 0.7 ml 0.5% Trypsin in PBS was added to the tube and incubated for 15 minutes at room temperature in a shaker at 400 rpm. Then, the supernatant was filtered through a cell strainer with mesh size 40 µm. 0.7 ml of Schneider’s S2 medium was added through the cell strainer and the 1.4 ml were transferred from a falcon tube to a fresh 1.5 ml Eppendorf tube. The filtrate was centrifuged for 7 minutes at 5000 rcf, washed with fresh 1.4 ml S2 medium and centrifuged again for 7 minutes at 5000 rcf. The pellet contained the obtained cells. The remaining tissue was digested again for 15 minutes with fresh 0.7 ml of 0.5% Trypsin in PBS. This process was repeated twice. The final pellet was resuspended in 20 µl fresh S2 medium and subsequently used for plunge freezing.

Plunge freezing of cells was performed on R2/1 SiO_2_ copper 200 mesh (Quantifoil) grids, glow discharged for 30 seconds. 4 µl of cell suspension were applied and plunge-frozen using a Vitrobot Mark 4 system (Thermo Fisher Scientific) at blot force 10, blotting time 10 s with blotting paper brought into contact with the grid on the backside only and Teflon sheets on both sides.

### Follicle cell identification on-grid and lamella preparation

After plunging, grids were assembled into AutoGrid cartridges and imaged on a Leica cryo-confocal microscopes based on the Leica TCS SP8 system (Leica Microsystems), equipped with a cryo-stage [79], 50x objective, NA 0.90, and two HyD detectors. A gridmap was acquired to identify suitable positions with fluorescent cells centered within grid squares. Fluorescence stacks were collected per grid squares at a zoom of 2, pinhole of 1 Airy Unit (AU), voxel size of 108 nm x 108 nm x 500 nm and 30 slices per position. The excitation laser has its emission peak at 488 nm and was used at 4% intensity. Fluorescence was detected using a HyD detector with its range set up from 495 nm to 541 nm at a gain of 100. Transmission was detected using a PMT detector at a gain of 560.

The grids were subsequently loaded into an Aquilos 1 cryo-FIB (Thermo Fisher Scientific) and overview SEM images were collected. EosFP-positive cells were identified based on the previous cryo-FLM experiments. On-grid lamellae were prepared at a stage tilt of 17° with a 45° pre-tilted shuttle (Thermo Fisher Scientific) resulting in a +-10° pre-tilt in the TEM similar to published protocols [22, 38]. Targeting of the nucleus was achieved using SEM imaging at 3 kV, 13 pA, 20 line integration at 100 ns dwell time, pixel width of 4-8 nm, and an image dimension of 3072x2188 pixels. Once a suitable nucleus position was found, circa 100-200 nm were removed from the top of the lamella to avoid SEM damage in the subsequent cryo-ET experiments and the lamella was thinned until contrast faded at 3 kV, 13 pA, 1 line integration at 200 ns dwell time and 1536x1094 image dimension.

### Cryo-lift-out from high-pressure frozen *D. melanogaster* egg chambers

Lift-out experiments were performed similar to previously described workflows [38]. 24 hours prior to dissection of egg chambers, the flies were transferred into fresh vials supplemented with yeast paste. Dissected egg chambers were high-pressure frozen in gold-coated copper Type A carriers (Engineering office M. Wohlwend) using the HPM010 (Wohlwend) or the EM Ice high-pressure freezing systems (Leica Microsystems). The egg chambers were added into the 100 µm or 200 µm cavity of the planchette and 20% Ficoll (70 kDa) in Schneider’s medium was used for filling the carrier. After freezing, the planchettes were pre-trimmed in a cryo-microtome (EM UC6/FC6, Leica Microsystems) operated at −170 °C using a 45° trimming knife with 6° clearance angle.

For fluorescence imaging, the planchettes were cryo-glued to custom 3D printed plastic replicas of the Leica cryo-shuttle and cryo-fluorescence light microscopy was done on a Leica TCS SP8 as specified above.

For LifeAct::GFP samples, stacks were collected at a zoom of 1.3, pinhole of 2 AU, excitation laser 488 nm at 6% and 552 nm at 6% for GFP and autofluorescence, respectively, and a voxel size of 170 nm x 170 nm x 860 nm with 38 sections around the focus position. Signal was detected using a HyD detector from 496 nm to 535 nm, a PMT for reflection light at 544 nm to 561 nm, and another HyD detector for autofluorescence from 593 nm to 669 nm. HyD gain was 100 and PMT gain was 400.

For emGFP::Nup358 samples, stacks were collected at a zoom of 2.6, pinhole of 1 AU, excitation laser 488 nm at 5% and 552 nm at 6%, and a voxel size of 42 nm x 42 nm x 480 nm with 75 sections around the focus position. Signal was detected using a HyD detector from 500 nm to 536 nm, a PMT for reflective light at 544 nm to 561 nm, and another HyD detector for autofluorescence from 593 nm to 669 nm. HyD gain was 100 and PMT gain was 400.

The planchettes and a half-moon grid were transferred to a standard 45° pre-tilted Aquilos (Thermo Fisher Scientific) shuttle into the FIB-SEM instrument for lift-out experiments. Correlation with the acquired fluorescence images was done using 3DCT [80] and trench milling was performed manually or automated in SerialFIB [38]. The custom pattern file was chosen to create a roughly 20 µm x 20 µm block [25] that could be undercut and lifted from the HPF sample. Lift-out device used was a Kleindiek micromanipulator [26] equipped with a cryo-gripper or a cryo-needle [23]. After transfer via redeposition (needle) or physical gripping (cryo-gripper), the attachment to the receiver grid was achieved using redeposition with multiple patterns at 0.3 nA, regular cross section, single pass, and a size of 1 µm. Note that for needle experiments, no copper block adapter was used in the presented data but is highly recommended for increased stability of attachment [25]. The coarse milling and polishing of the lamellae were performed using an end-pointing approach with SEM data acquisition to target the follicle cell nuclear envelope. Approximately 4 µm of volume imaging was performed prior to lamella preparation at a section thickness of 100 nm and SEM imaging at 3 kV, 13 pA, 50 line integration at 200 ns dwell time, pixel width of 6.84 nm, and an image dimension of 3072x2188 pixels. Images were acquired with the Everhart-Thornley detector (ETD) in Standard mode or with optional addition of in column T1 and T2 detector in OptiTilt mode. Milling for volume imaging was performed at 30 kV 0.5 nA using regular cross-sections. Once the feature of interest, here the nuclear envelope, was visible in the SEM image, further imaging and milling was stopped, approximately 100 nm were removed from the top surface and the rest of the lamella preparation was performed from below with sequentially decreasing milling currents of 0.5 nA to 3 µm thickness, 0.3 nA to 1 µm, and 0.1 nA to 500 nm. Final polishing was done at 50 pA until the contrast fainted at 3 kV 13 pA SEM image. This yielded a lamella below 200 nm in thickness that was transferred to the TEM.

### Cryo-ET data acquisition

Cryo-ET data was acquired on a G3 Titan Krios equipped with a Falcon 4i and Selectris X energy filter. A dose-symmetric scheme was used with a 3° increment at a dose per tilt of 3 e/Å^2^ for on-grid lamellae and a 2° increment at a dose per tilt of 2 e/Å^2^ for lift-out lamellae. Total dose in both cases was 120 e^-^/Å^2^. Tilt range was from −50° to +70° or −70° to +50° depending on the loading direction with a pre-tilt of +10° and −10°, respectively. Tomograms were recorded at a nominal magnification of 42.000x resulting in a pixel size of 2.96 Å.

Data was pre-processed using the Tomoman version 0.7 pipeline (https://github.com/williamnwan/TOMOMAN). EER files were rendered as frames aiming at 0.2 e^-^/Å^2^ dose per frame and motion-corrected using the MotionCor2 version 1.4.7 [81]. The CTF was estimated with CTFFIND4 version 4.14 [82]. Bad tilts were removed manually using the dedicated Tomoman scripts. Dose weighting was performed in either Tomoman or Warp version 1.0.9 [83]. Tomograms for initial subtomogram averaging were reconstructed using automated reconstruction in AreTomo version 1.3.3 [84] or patch tracking in IMOD version 4.12.32 [76, 85] at pixel size of 11.84 Å using patches of 200x200 pixels and a fractional overlap of 0.45 in X and Y.

### Cryo-ET data analysis

#### Initial subtomogram average of entire VLPs

Initial template matching was done with Stopgap (0.7.0) [86] (pixel size 11.84 Å) using a hollow sphere with an outer diameter of 505 Å and an inner diameter of 249 Å as template. As the reference was a sphere, only a single angle was matched. The particle positions were extracted from the score map with a threshold adjusted per tomogram to avoid false positives, generally with cross correlation scores between 0.185 and 0.27. CTF fitting was done in Warp and 1780 particles were extracted in a box size of 192^3^ voxels and a voxel size of 2.96Å. To avoid alignment to the missing wedge random Euler angles were assigned to the particles. Next, we performed a single class alignment in Relion version 3.0 without symmetry and sphere with a diameter of 505 Å as reference. The angular search was restricted to 36° with an increment of 3.7°. The resulting structure showed clear six-fold and five-fold positions, suggesting an icosahedral lattice. To generate a reference for refinement, the C1 reference was aligned to an I1 symmetric pose and symmetry was applied using relion_image_handler. Finally, a refinement with I1 symmetry and full angular search with an increment of 3.7° was performed.

#### Refinement and Classification of VLPs

Particles were re-extracted in Warp (pixel size 2.54Å, box of 232 voxel) and aligned in Relion 3.0 using the scaled reference obtained from the step before. A soft mask around the capsid was applied. Particle Poses, Stage Angles and Deformation were refined in M version 1.0.9 [87]. A second round of M refinement with additional Defocus processing was carried out. Particles were re-extracted in M with a pixel size of 2.96Å and a Box of 192^3^ voxels. A 3D classification with 6 classes was performed in Relion (local Search of 1.8° and increment of 0.9°). 2 classes turned out to be empty. The best resolved class (735 particles) of the remaining four was chosen for refinement. Subsequently, the particles from this class were re-extracted in M with a pixel size of 2.54Å and a box size of 256^3^ Voxels and refined in Relion. In the final refinement, a mask around the capsid protein was applied that removed the inner density annotated as ribonucleoprotein from alignment. Post-processing was carried out in Relion, resulting in a resolution of 7.7 Å measured by gold-standard Fourier shell correlation (GSFSC).

### Model building of the *copia* pentameric and hexameric structures

Based on the 7.7 Å reconstruction of the *copia* capsid, models for the *copia* capsid were created starting off with the pentameric C5 and hexameric C3 environments of the structure. Five and six copy assemblies of the N-terminal 270 residues of the *copia* Gag protein were predicted using AlphaFold-Multimer version 2.2.0 [88] and rigid body fitted into the corresponding electron microscopy map. While the structure of six monomers fitted well into the C3 density, the five-fold symmetric map showed a discrepancy between the prediction and the reconstructed C5 map. The C-terminal domain needed to shift slightly in order to make the secondary structure elements of prediction fit the cryo-EM map. The structures were fitted in ISOLDE version 1.5 [89] within ChimeraX version 1.5 [90]. Simulations were run at 100 K using the AMBER ff14sb force field [91]. The solvent is treated using GB-Neck2 implicit solvent [92]. Additionally, grid-based protein backbone corrections were used. The entire capsid structure was assembled from these subunits using BioMT matrices for icosahedral I1 symmetry.

### Cn-Positions (C1, C3, C5) and contacts on the Bravais lattice

To address lattice defects, competitive template matching on individual *copia* particles was performed. M was used to generate deformation corrected full *copia* particles (box 272x272x272 voxel size 2.96Å) from the M run that yielded the 7.7 Å capsid structure. Three templates were generated from the refined full *copia* capsid average by cropping subvolumes around the C1, C3 and C5 positions (compare Fig. 3D) with a box size of 128x128x128 pixels and a pixel size of 2.96 Å. For each template, an adapted version pyTOM-template matching [93] was run with an angular increment of 7°. PyTOM-tm was adapted to use the point spread function of M corresponding to each particle. The cross-correlation scores from each template (c1, c3, c5) were combined by a maximum operation. Template numbers and angles were bookkept in separate volumes according to their maximum score. Next, 92 peaks were extracted from the combined cross correlation function using custom MATLAB scripts based on the tom-toolbox [94]. Corresponding angles and template numbers were determined. Positions were filtered by cross-correlation score (> 0.1), radius (< 272 Å) and normal angle deviation (< 40°). Locations were transformed back to their absolute position in the tomogram by adding the corresponding center of the *copia* particle to the extracted Cn-Positions. This resulted in 117,963 positions (C1: 68,190, C3: 31,563, C5: 18,210). Finally, the subvolumes of the corresponding locations were reconstructed in Warp with a box 168x168x168 of and a voxel size of 2.24 Å.

Each Cn-position was shifted to the edge of the particle. Nearest neighbor distance was determined for each Cn-position and shown as a histogram. Suggested by the first minima of the distance histogram, Cn-positions with a spacing smaller than 36 Å were classified as contacts. The latter procedure identified 5620 contact positions. Next these contact points were grouped according to their Cn-position pairing (c1-c1, c1-c3, c1-c5 …). For each Cn-position pairing an average was created using relion_reconstruct (filtered to 35 Å). The random occupancy of each Cn-position pairing was determined by selecting a random subset of the original Cn-Position classes with the same amount as the contact points. Then, class pairs were generated and counted. This was repeated 1000 times. Mean and standard deviation were calculated for each of the classes. The relative measured abundance combined with the random relative abundance was shown in a histogram. Three standard deviations were used as error bars for the random abundance.

### *Copia* inter particle lattice

To determine the *copia* lattice organization, particle centers from an area with a constant grid were used (pixel size of coordinates: 5.92 Å). For each position, the neighboring positions within a radius of 600 Å were determined. Neighboring locations were bookkept in a volume with 200x200x200 pixels. Finally neighboring positions volumes were summed to obtain the neighbor map, which can be interpreted as a 3D histogram. The same procedure was performed for contact positions (C1-C5, C1-C1 and C5-C5). This was done with an adapted version of STOPGAP’s [86] sg_neighbor_plot_local. The function was modified to ignore the pose of the particles. Next the 12 highest peaks were extracted from the neighbor map. Vectors between the center and the peak positions were calculated. Three non-symmetric vectors were chosen to describe the Bravais lattice. The length and the angle between the vectors were determined to characterize the lattice. (Length: 528 Å, 518 Å, and 522 Å; Angle: 62.0°, 61.3°, and 61.2°). For visualization, the 3 grid vectors were plotted for each *copia* position to resemble the grid. To display the *copia* connections 2 additional vectors were extracted and displayed. Contact position peaks were used to assign a class to particle center vectors.

### Subtomogram averaging of NPCs for tomographic backmapping

The nuclear pore complex subtomogram averaging was performed as described previously [95]. In brief, particles were picked manually in IMOD using custom scripts developed by Dr. Beata Turonova (MPI Biophysics, Frankfurt). Subtomogram averaging was performed in tomSA. Manually picked particles were aligned at a pixel size of 23.68 Å (bin8) in a box of 72^3^ voxels with C8 symmetry applied. The alignment was refined at 11.84 Å (bin4). As the particle number was limited (36 NPCs), entire pores were averaged with applied symmetry and not split into asymmetric subunits.

### Template matching of ribosomes for tomographic backmapping

PyTom-tm template matching was performed based on the *D. melanogaster* 80S ribosome structure (PDB ID: 4V6W) as template. The angle increment was 11° and the template was filtered to 25 Å resolution. 897 particles were extracted using pytom_extract_candidates.py with Number of false positives set to 1. Subtomograms were averaged in STOPGAP v0.7 at bin4, corresponding to a pixel size of 11.84 Å and backmapped to the tomogram for visualization in ArtiaX version 0.1 [96]. Membrane segmentation was performed using Membrain-Seg version 9 [97].

### Radial averages and VLP diameters

3D volumes of *copia* VLPs in a box size of 64^3^ voxels at 4 times binning corresponding to a pixel size of 11.84 Å were extracted for radial averaging. Radial average profiles were calculated from the extracted subvolumes using e2proc3d.py in EMAN2 [98], averaged in numpy [99], and plotted using matplotlib. Statistical significance of VLP sizes for single comparisons was determined using the unpaired Student’s t test. Ns: p > 0.05. *: p ≤ 0.05, **: p ≤ 0.01, ***: p ≤ 0.001, ****: p ≤ 0.0001

### Visualizations

Data was visualized in IMOD version 4.12.32 [76] and FIJI [100]. 3D renderings were done in ChimeraX version 1.5 [90] using the ArtiaX plugin version 0.3 [96].

### Key resource table

| Reagent type (species) or resource | Designation | Source or reference | Identifiers | Additional Information |
| --- | --- | --- | --- | --- |
| Genetic reagent (*Drosophila melanogaster*) | UAS-Sec31::RFP | Blooming ton<br>Drosophila<br>Stock Center | FlyBase ID:<br>FBst0086533 | w* P{UAS-Sec31.RFP}13.5; snaSco/CyO, P{ftz-lacB}E3 |
| Genetic reagent (*Drosophila melanogaster*) | GFP::nup358 | (Hampoelz et al., 2019) | - | emGFP::nup358 PBac{y[+mDint2] = vas-Cas9}VK00027 |
| Genetic reagent (*Drosophila melanogaster*) | Tj-gal4 | (Cassella & Ephrussi, 2022);<br>Gift of Dr. Anne Ephrussi | - | ;tj-Gal4/CyO;; |
| Genetic reagent (*Drosophila melanogaster*) | UASp-Sec13::EosFP | generated by Dr. JM Philippe and Dr. Bernhard Hampoelz, provided by BH | - | w* P{UASp-Sec13::EosFP}/ CyO ;) |
| Genetic reagent (*Drosophila melanogaster*) | GFP-Nup107 | Blooming ton<br>Drosophila<br>Stock Center | FlyBase ID:<br>FBst0035514 | w*; P{GFP-Nup107.K}9.1 (BL-35514) |
| Genetic reagent (*Drosophila melanogaster*) | LifeAct::GFP | Provided by Dr. Cristina Pallares Cartes | - | w*; P{w+, sqhp>LifeAct::mNeon Green}; MKRS/TM6. |
| Genetic reagent (*Drosophila melanogaster*) | *copia* dsh TL63 | This study | - | w; *copia* dsh attP2 (TL63) |
| Genetic reagent (*Drosophila melanogaster*) | *copia* sh TL67 | This study | - | w; *copia* sh attP2 (TL67) |
| Genetic reagent (*Drosophila melanogaster*) | *copia* sh TL68 | This study |  | w; *copia* dsh attP2 (TL68) |
| Antibody | *Copia* Polyclonal antibody | Cusabio | CSB-PA366106XA01DLU |  |
| Antibody | rabbit anti-McClintock Gag polyclonal Antibody raised against peptide ITEAQTAEFRPQASE QANS | (Senti et al., 2023) |  |  |
| Antibody | mouse anti-Aub monoclonal Ab 8A3-D7 | (Senti et al., 2015) |  |  |
| Antibody | rabbit anti-Ago3 polyclonal Ab | (Mohn et al., 2014) |  |  |
| Antibody | mouse anti-laminDM0 monoclonal ADL84.12 |  |  |  |
| Reagent | *copia* smFISH probe set | - | this study | Probe sequences in supplement |
| Reagent | Maverick smFISH probe set |  | this study | Probe sequences in supplement |
| Software, algorithm | SerialFIB | https://github.com/sklumpe/SerialFIB (Klumpe, 2021) |  |  |
| Software, algorithm | AutoScript4 | Thermo Fisher Scientific |  |  |
| Software, algorithm | LAS X | Leica Microsystems |  |  |
| Software, algorithm | Fiji | https://imagej.net/software/fiji/ |  |  |
| Software, algorithm | 3DCT | https://3dct.semper.space/index.html |  |  |
| Software, algorithm | SerialEM | https://bio3d.colorado.edu/SerialEM/ |  |  |
| Software, algorithm | Warp | http://www.warpem.com/warp/ |  |  |
| Software, algorithm | TOMOMAN | https://github.com/williamnwan/TOMOMAN |  |  |
| Software, algorithm | MotionCorr2 | https://emcore.ucsf.edu/ucsf-software |  |  |
| Software, algorithm | NovaCTF | Turoňová et al., 2017b |  |  |
| Software, algorithm | CTFFIND4 | https://grigoriefflab.umassmed.edu/ctffind4 |  |  |
| Software, algorithm | IMOD | https://bio3d.colorado.edu/imod |  |  |
| Software, algorithm | pyTOM | https://pytom.sites.uu.nl/ |  |  |
| Software, algorithm | RELION | https://www3.mrc-lmb.cam.ac.uk/relion/index.php/Main_Page |  |  |
| Software, algorithm | STOPGAP | Wan, 2020 |  |  |
| Software, algorithm | UCSF ChimeraX | https://www.rbvi.ucsf.edu/chimerax/ |  |  |
| Software, algorithm | Membrain-seg | https://github.com/teamtomo/membrain-seg |  |  |
| Software, algorithm | ISOLDE | https://tristanic.github.io/isolde/ |  |  |
| Software, algorithm | Seurat | https://satijalab.org/seurat/ |  |  |
| Software, algorithm | SoloTE | https://github.com/bvaldebenitom/SoloTE |  |  |
| Dataset | scRNA-seq *D. melanogaster* ovary | Jevitt A et al. PLoS Biology (2020) | SRA ID: SRX7814226<br>GEO ID: GSE146040 |  |

### SmFISH probe sequences

See attached source files.

