## Supplementary Information for "In-cell structure and snapshots of *copia* retrotransposons in intact tissue by cryo-electron tomography"

**Movie S1: Tour of the in-cell *copia* structure, related to Figure 2.** The movie builds up the *copia* capsid structure from the asymmetric unit containing the two C1 conformations colored in the same color for simplicity (yellow), three copies of the C3 conformation 1 (blue), three copies of the C3 conformation 2 (red) and one copy of the C5 conformation (orange). The movie goes on to show the assembly of the entire icosahedral lattice based on this asymmetric unit. Subsequently, the coloring is changed to a radial color scheme and the structure is cut open to reveal the ribonucleoprotein density at the luminal side (red) engulfed by capsid (blue and orange). The movie ends by morphing into the structural model of the *copia* capsid.

**Movie S2: Tomogram and 3D rendering of nuclear *copia* cluster.** The movie starts by moving back and forth through a tomogram of large nuclear *copia* clusters collected from isolated follicle cells. The 3D rendering of particle positions from *copia* subtomogram averaging is subsequently revealed.

**Movie S3: Visualization of the organization within nuclear *copia* cluster.** First, the movie zooms in on a patch of *copia* VLPs that is subsequently extended for visualization purposes. The full capsid structure is then faded to a silhouette representation and particle centers are displayed by yellow spheres. The intercapsid contacts are visualized between these spheres for C5-C5 (red), C1-C5 (yellow), and C1-C1 (blue) contacts.

**Movie S4: Tomogram, 3D rendering and animation of *copia* VLPs in vicinity of nuclear pores from intact *D. melanogaster* egg chambers.** First, the movie goes back and forth through the tomogram collected on a lift-out lamella from the follicle cell epithelium of intact *D. melanogaster* egg chambers fixated by high-pressure freezing. Subsequently, the segmentation of microtubules (cyan), membranes (grey), ribosomes (white), cytoplasmic VLPs (orange), nuclear VLPs (yellow) and nuclear pore complexes (purple) is revealed. A zoom onto the cytoplasmic VLPs interacting with the nuclear pore and a perspective through the nuclear pore complex is given, illustrating the size difference of cytoplasmic and nuclear VLPs.

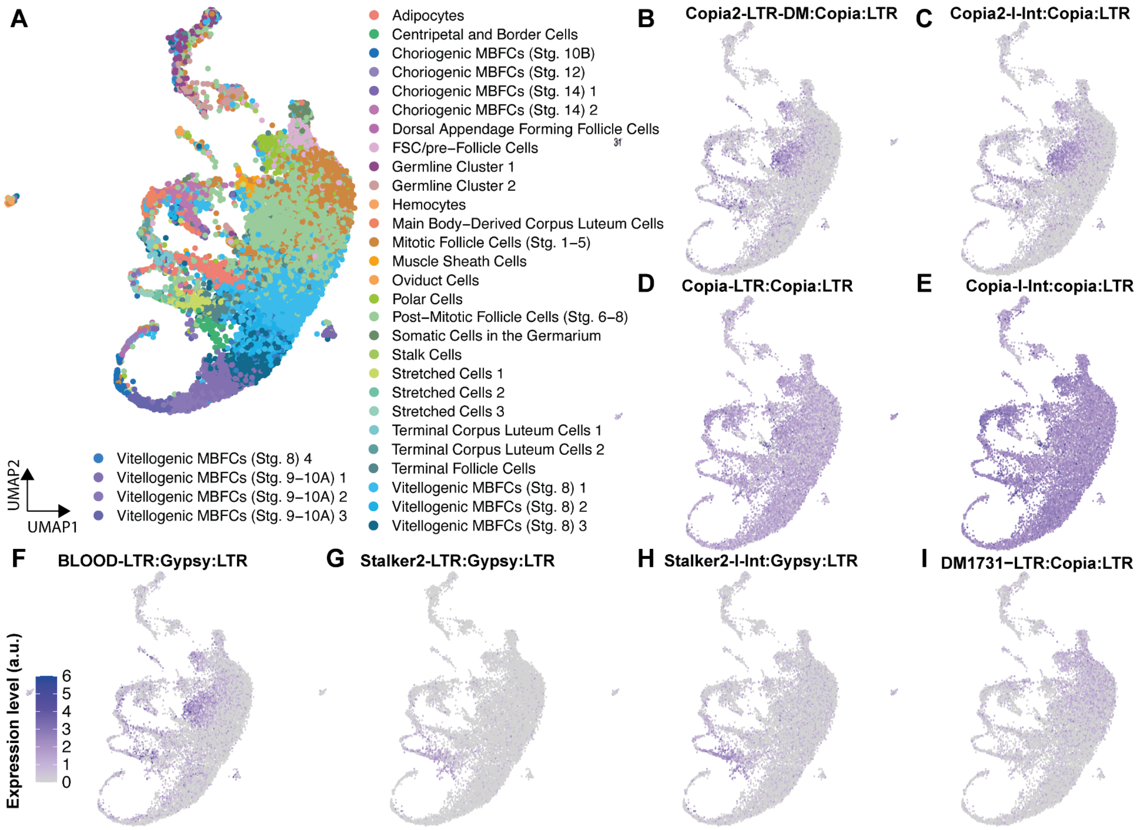


**Fig. S1: Ovary scRNA-seq analysis of *copia* and other LTR- elements. A:** UMAP clustering including labels for clusters using cell markers as performed previously [1]**.** The determined clusters from re-analyzed raw data fitted the previous labels and, thus, were used to describe expression level of the respective genes (see Materials & Methods). This allowed for the identification of LTR elements that are indicated to be specifically expressed in certain cell types within the *D. melanogaster* ovary tissue. **B-I:** Examples of expression level of a number of LTR-elements present in the dataset. Note the absence of signal within the follicle cell epithelium (main body follicular cells, MBFC, and mitotic follicle cells in the mapping shown in **A** for most elements. *Copia*, on the other hand, gives clear signal throughout somatic cells **(D-E)**. Analyzed were w[-] flies (BL#3605) [1].

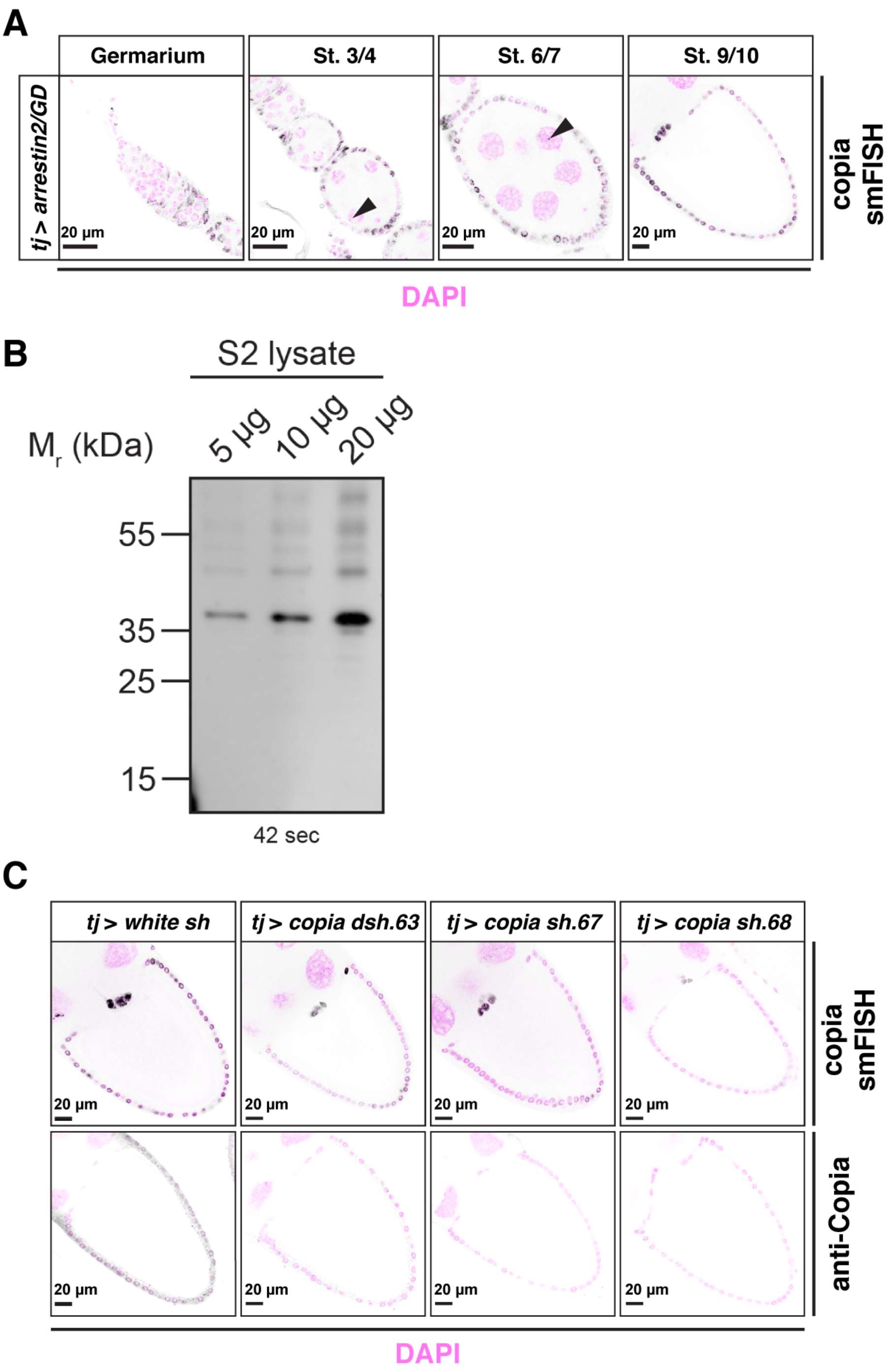


**Fig. S2: *Copia* antibody validation and broad *copia* expression across somatic follicle cell stages. A:** Confocal images of *copia* smFISH (black) and DAPI (magenta) throughout the different oogenesis stages of control knockdowns (*tj*>*arrestin2/GD*) shows widespread expression in the ovarian somatic cells. Asterisks indicate nuclear punctae likely representing nascent *copia* piRNA precursor transcripts **B:** Western blot analysis of *copia* antibody against different amounts of S2 cell lysate. **C:** Confocal micrographs of *copia* smFISH (top panel) in black and anti-*copia* (Gag) staining (bottom panel) in black and DAPI (magenta) in stage 10 ovary of control knockdown (*tj*>*white* sh) and three different *copia sh* knockdown genotypes (*tj*>*copia* sh, see Mat&Methods). Note the decrease of signal in the ovarian somatic cells in both the smFISH and anti-*copia* immunofluorescence. Arrowheads indicate discrete foci in nurse cell nuclei, likely representing nascent *copia* piRNA precursor transcripts.

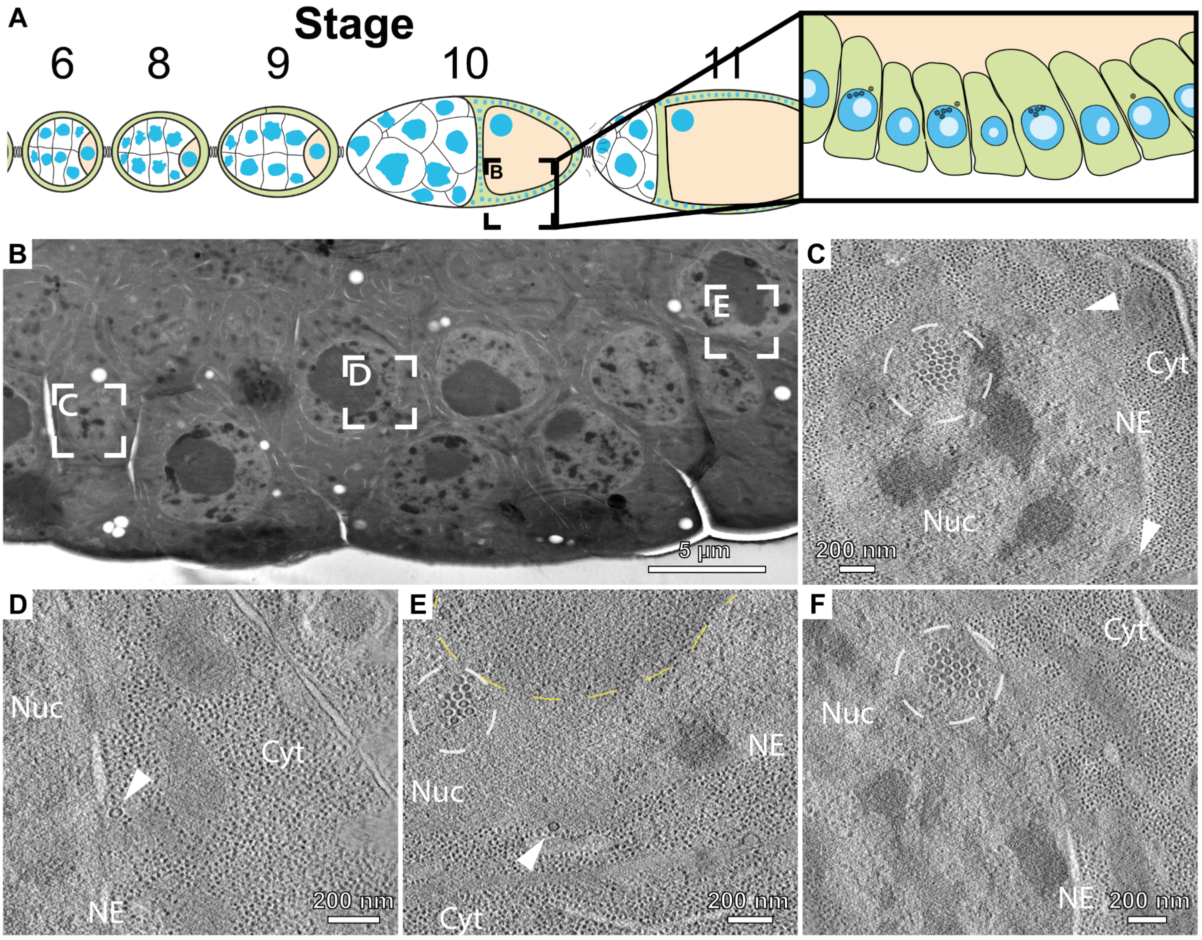


**Fig. S3: Electron tomography on thin sections of plastic-embedded egg chambers. Additional examples from the section presented in Figure 1. A**: Schematic of the origin of the section. The follicle cell epithelium of stage 10 egg chambers was targeted. **B**: Overview of the plastic section. Insets represent the location of the tomograms shown in C-E. **C-F**: Slices through tomograms collected at the nuclear periphery of the follicle cell nuclei. Marked are VLP clusters (white dashed circles), cytoplasmic VLPs (arrowheads), the nuclear envelope (NE), cytoplasm (Cyt), nucleoplasm (Nuc), and the nucleolus (yellow dashed circles). Tomogram from F originated from a different section than depicted in B. Genotype of used flies was *tj>Sec31::mCherry*.

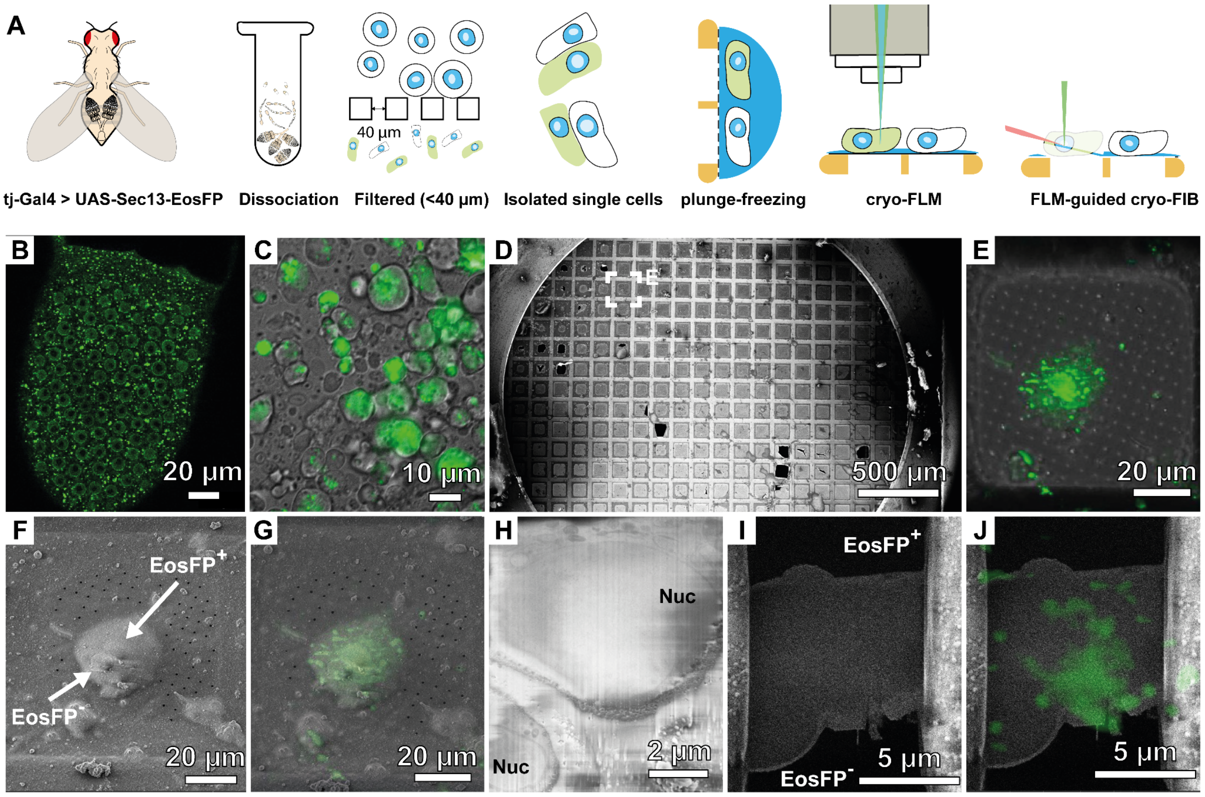


**Fig. S4: Isolation protocol for on-grid lamella preparation of tissue-specific cells using the UAS-GAL4 system. A:** Schematic workflow of the isolation and on-grid identification procedure of Sec13::EosFP positive cells from *tj>Sec13::EosFP* flies. **B:** Representative image of a stage 10 egg chamber of *tj>Sec13::EosFP* flies. **C:** Image of cells used for plunge freezing after tissue dissociation and filtering. **D:** SEM overview of a grid prepared from the isolated cells. Rectangle indicates one of the positions that were subjected to cryo-FIB milling. Grid square shown in the subsequent panels is indicated by a rectangle. **E:** Maximum intensity projection (MIP) of the cryo-confocal fluorescence light microscopy stack collected on the grid square indicated in D. **F:** SEM image of E shows two cells adjacent to each other, one positive and one negative for EosFP signal. **G:** Overlay of MIP from E and SEM image from F. **H:** Secondary electron SEM image of the surface after ablation of approximately 2 µm of cellular material from the top using the Everhart Thornley detector. Clearly visible are the two nuclei (Nuc). **I:** SEM image of the final thinned lamella. **J:** Overlay of FLM image with the final lamella image from I.

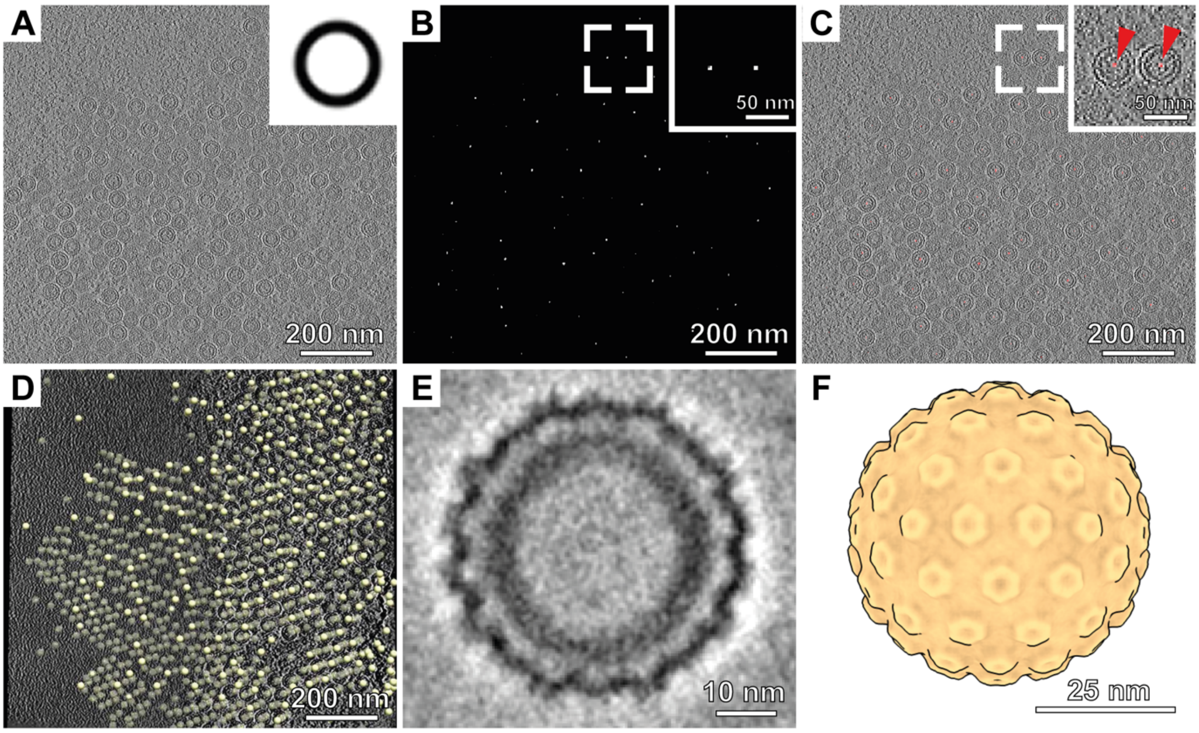


**Fig. S5: Initial average of *copia* VLPs from nuclear clusters via template matching with a hollow sphere and C1 alignment. A:** Tomographic slice from a reconstructed tilt series collected on a nuclear *copia* VLP cluster. Inset shows the reference used for template matching. **B:** Scoring map of template matching with the hollow sphere depicted in A. Inset shows zoom on two VLP peaks. **C:** Overlay of the tomographic slice (gray) and scoring map (red) from A-B. Inset shows the same region as in the inset in B. Red arrows indicate template matching peaks **D:** Backmapping of the identified *copia* particles from template matching into the 3D volume of the reconstructed tomogram. **E:** C1 reconstruction of the particles from D. **F:** Render of the volume depicted in E displayed with applied icosahedral symmetry I1.

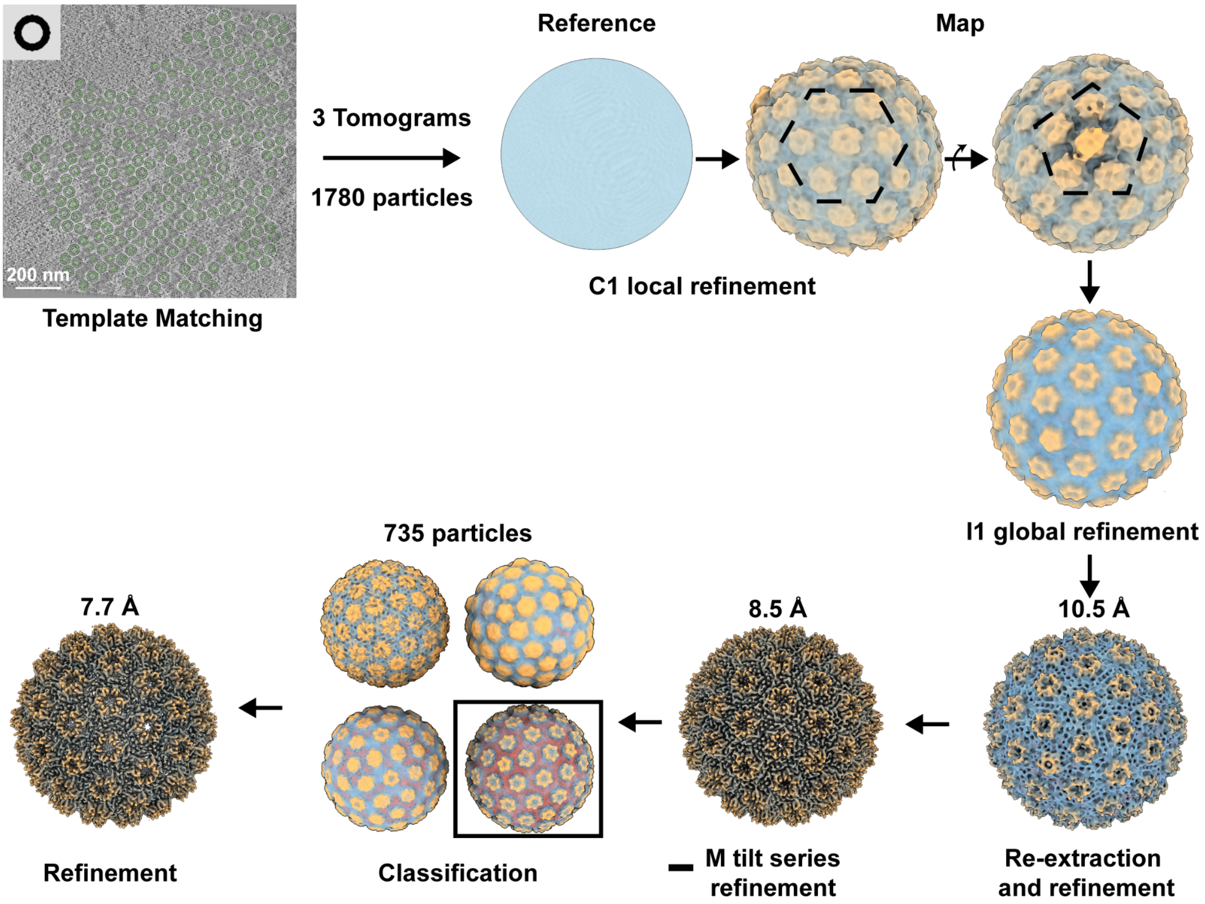


**Fig. S6**: Subtomogram averaging workflow using Warp/Relion3.1/M for full capsids. C1 refinement of 1780 particles obtained by template matching using a spherical reference resulted in a map that already shows the typical C5 and non-symmetric C6 geometries of icosahedral assemblies. Based on that observation, icosahedral symmetry was used in the subsequent steps. I1 global refinement and subsequent re-extraction and refinement yielded a structure at 10.5 Å resolution. After M tilt series refinement, the resolution reached the subnanometer regime at 8.5 Å. Subsequent 3D classification cut down the particle number to 735 particles, removing particles that are likely damaged or cut at the lamella edges. The final structure after refinement gave a resolution of 7.7 Å.

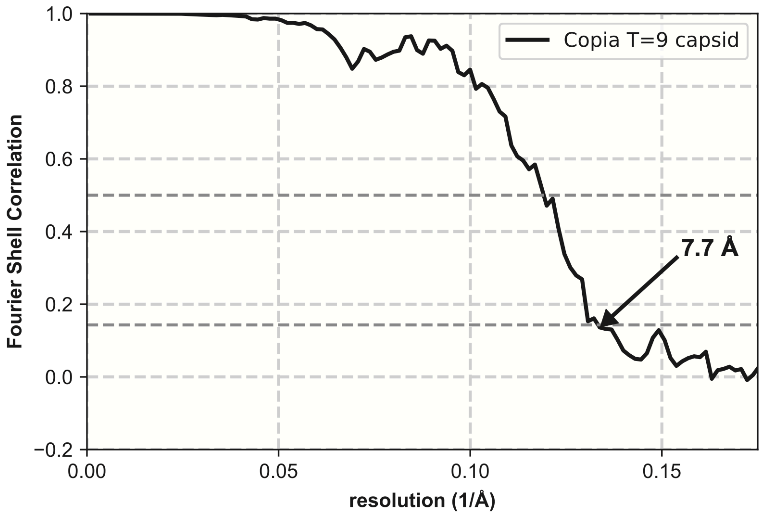


**Fig. S7**: Gold-standard Fourier shell correlation (GSFSC) curve for final structure of T=9 *copia* capsid. Cutoffs shown are at 0.5 and 0.143 FSC (dashed line). Reported resolution of 7.7 Å at 0.143 FSC.

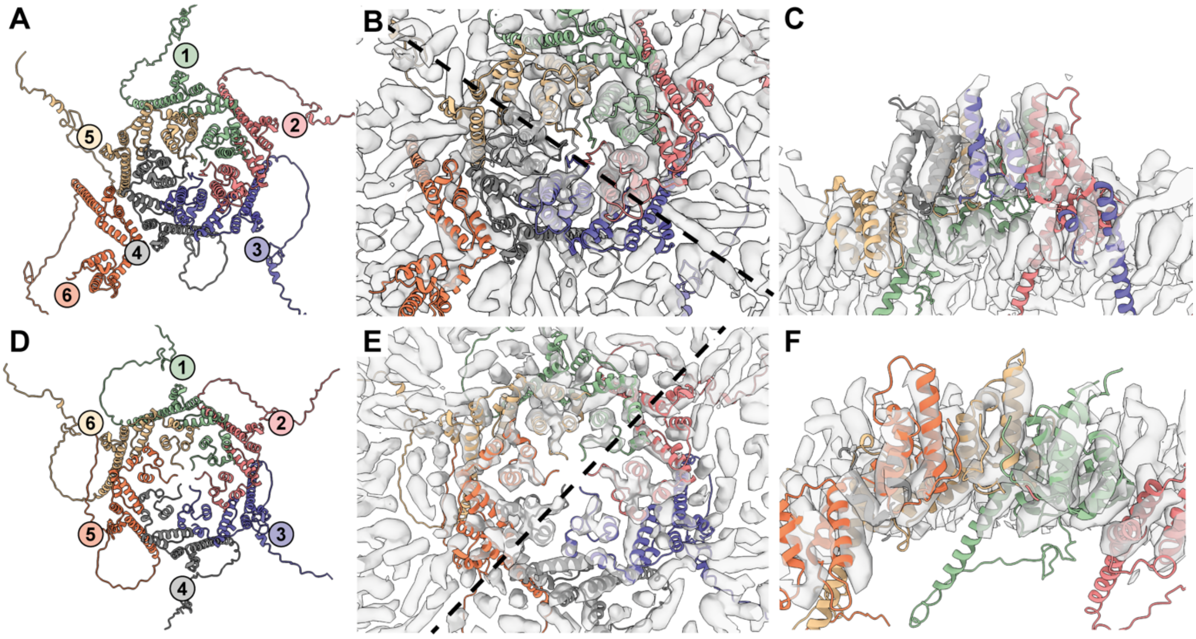


**Fig. S8: AlphaFold-multimer predictions yield pentameric and hexameric structures. A:** AlphaFold-multimer prediction for 6 *copia* Gag monomers (residues 1-270) yielding a five-fold symmetric structure. **B:** Rigid body fit of the 5-fold symmetric structure from A show good fit of the NTD domain (center) but no helix correspondence of the CTD domain. Dashed line indicates the plan of the side-view in C. **C:** Side-view of the rigid-body fit of the pentameric structure as indicated in B. **D:** AlphaFold-multimer prediction for 6 *copia* Gag monomers (residues 1-270) yielding a six-fold symmetric structure. **E:** Rigid body fit of the prediction in D into density of the non-symmetric hexamer from the icosahedral structure. Note that the helices of the CTD have corresponding density in this hexameric assembly. Dashed line indicates the plane of the side-view in F. **F:** Side-view of the rigid-body fit of the hexameric structure as indicated in E.

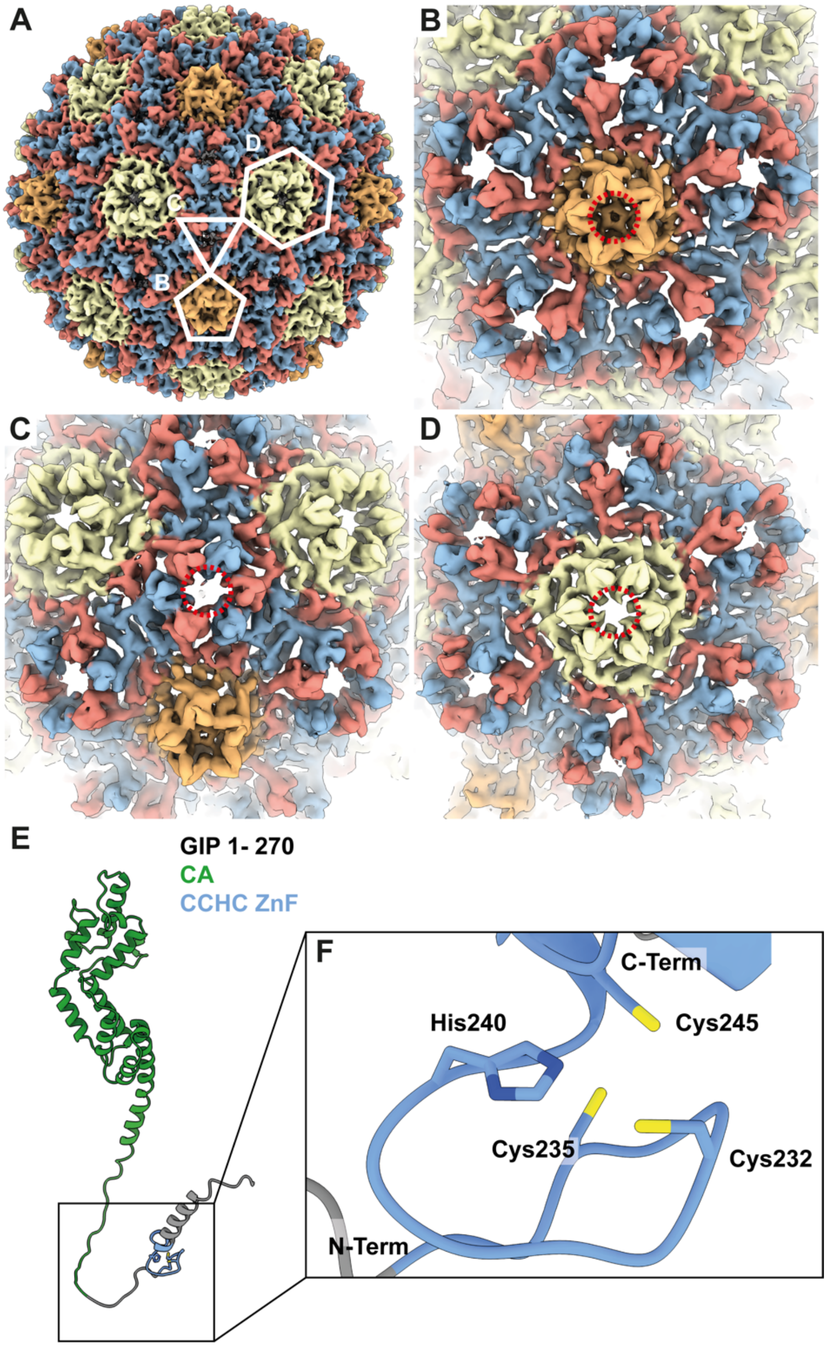


**Fig. S9: Immature *copia* capsid show pores in hexameric but not pentameric assemblies and AlphaFold prediction of full-length *copia* Gag-Int-Pol (GIP) show zinc finger (ZnF) module. A:** Overview image of the *copia* capsid structure. Pentagon, triangle, and hexagon show the positions of zoom images depicted in B, C, and D, respectively. **B:** Zoom on a C5 environment. The red dashed circle indicated the density present closing the pore in the capsid lattice. **C:** Zoom on a C1 environment. Note the missing density within the lattice pore, indicated by a dashed red circle. **D:** Zoom on a C3 environment. Dashed red circle indicates the pore in the capsid lattice. All images in A-D kept at the same threshold. **E:** AlphaFold prediction of residues 1 to 270 from full length *copia* Gag-Int-Pol (GIP) (EBI AlphaFold Database, UniProt ID: P04146) with NTD and CTD of CA in green and CCHC ZnF site in lightblue. **F:** CCHC ZnF formed by residues His240, Cys 235, Cys 232, Cys245, predicted by ProSite as a zinc finger module.

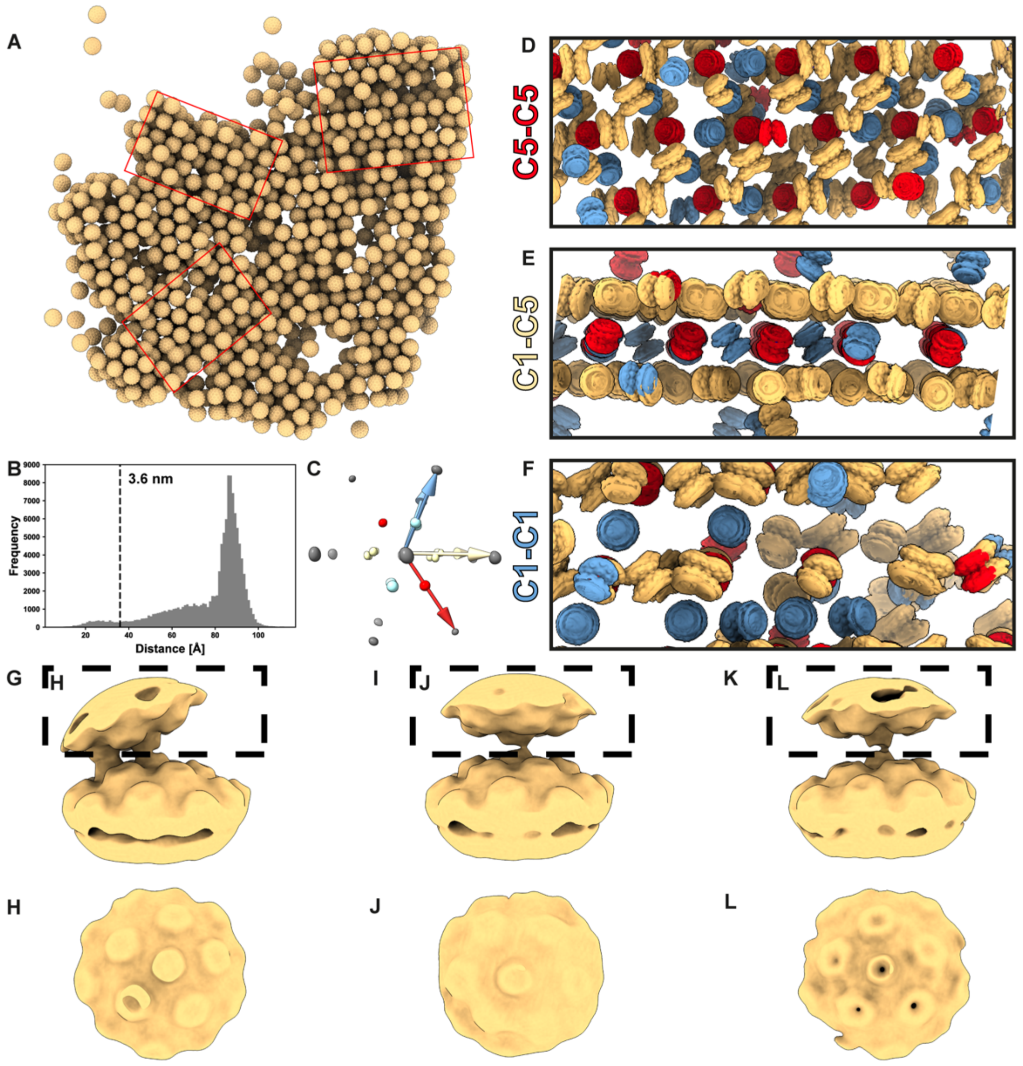


**Fig. S10: Intercapsid contacts form regular arrays. A:** 3D rendering of capsid positions from the tomogram shown in Figure 2 demonstrates the high mosaicity (red rectangles) of the lattice within nuclear clusters. **B:** Nearest neighbor distance determined for each Cn-position (C1, C3, C5) plotted as a histogram. Suggested by the first minima of the distance histogram, Cn-positions with a nearest neighbor distance smaller than 36 Å were classified as contacts. This procedure identified in total 5620 contact positions. The highest peak at 9 nm represents the neighboring intra-particle position on the icosahedral surface of the individual VLP. **C:** 3D Histogram of neighboring VLPs within nuclear *copia* clusters. The three main capsid-capsid interactions C1-C1 (blue), C5-C5 (red), and C1-C5 (yellow) are displayed as colored arrows within the 3D histogram. **D-F:** Backmapping of the contact subtomogram averages to their respective particle positions. The placed particles are shown from different angles along the **D:** C5-C5 (red), **E:** C1-C5 (flax), and **F:** C1-C1 direction (blue). Note the regularity in contact patches. **G:** Side view of the C1-C5 contact map and **H:** top view of the upper contacting capsid density depicted in G (black dashed rectangle). **I:** Side view of the C1-C1 contact map and **J:** top view of the upper contacting capsid density depicted in I (black dashed rectangle). **K:** Side view of the C5-C5 contact map and **L:** top view of the upper contacting capsid density depicted in K (black dashed rectangle).

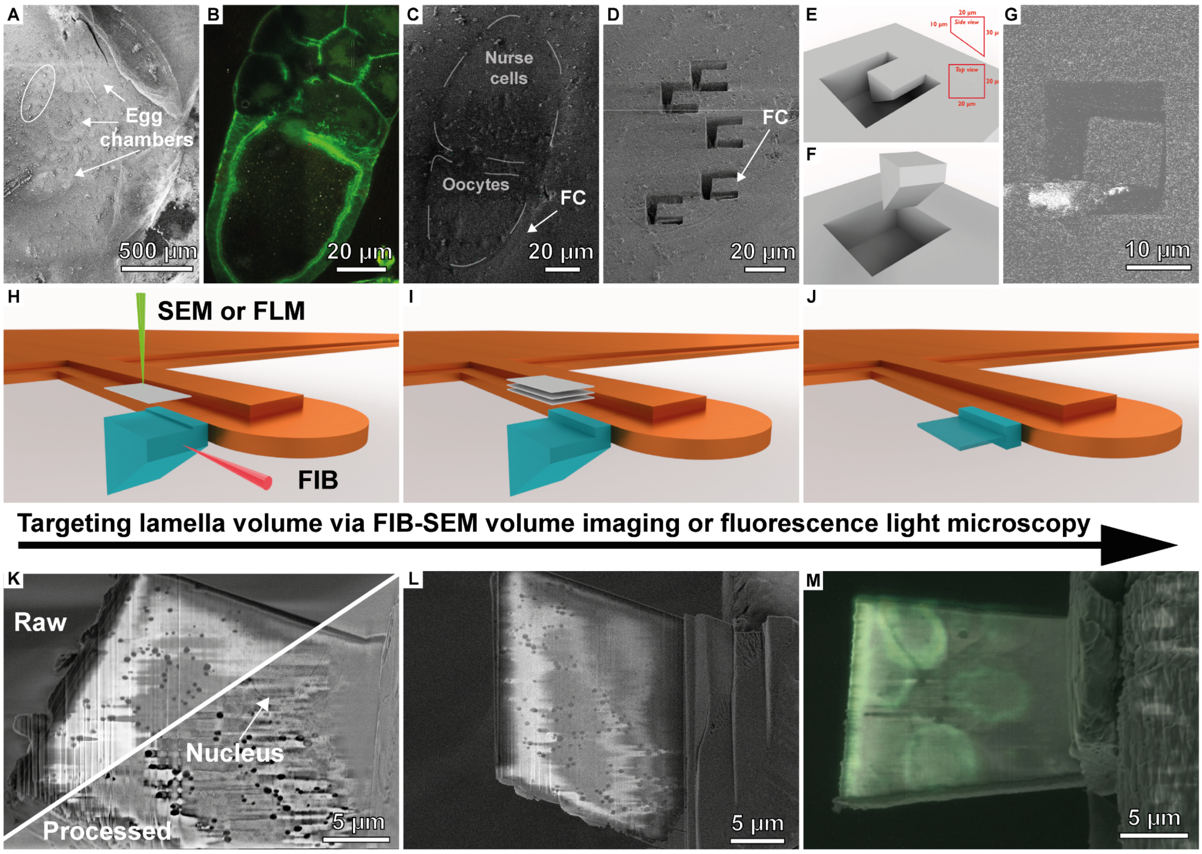


**Fig. S11:** Lift-out experiments and targeting in *D. melanogaster* egg chamber. **A:** Overview SEM image of the high-pressure freezing planchettes. The egg chamber used in the subsequent lift-out experiment is indicated by a white circle. **B:** Fluorescence and **C:** FIB image of the egg chamber, indicating major features such as nurse cells, oocytes and follicle cells (FC). **D:** SEM image after trench milling. **E-F:** Geometries of the lift-out volumes. **G:** Image of attachment of a cryo-needle during lift-out. **H:** Schematic showing targeting possibilities in cryo-lift-out. Either scanning electron microscopy (SEM) or fluorescence light microscopy (FLM) can be used to target specific events. **I:** When targeting using FIB-SEM volume imaging, SEM images are obtained from the surface of the lift-out volume until the feature of interest is seen and a lamella is thinned from the bottom. **J:** Lamella thinned after targeting. **K:** SEM image of the surface of the lift-out volume showing the targeted follicle cell nuclei in the raw (Raw) and post-processed (Processed) image. **L:** Overview image of the final thinned lamella from K. **M:** Overview image of the final lamella image of a different experiment. Here, integrated fluorescence light microscopes were used for targeting and an overlay of the SEM image and the *GFP::Nup358* green fluorescence is shown. Genotypes: **A-L** *LifeAct::GFP*, **M**: *GFP::Nup358*

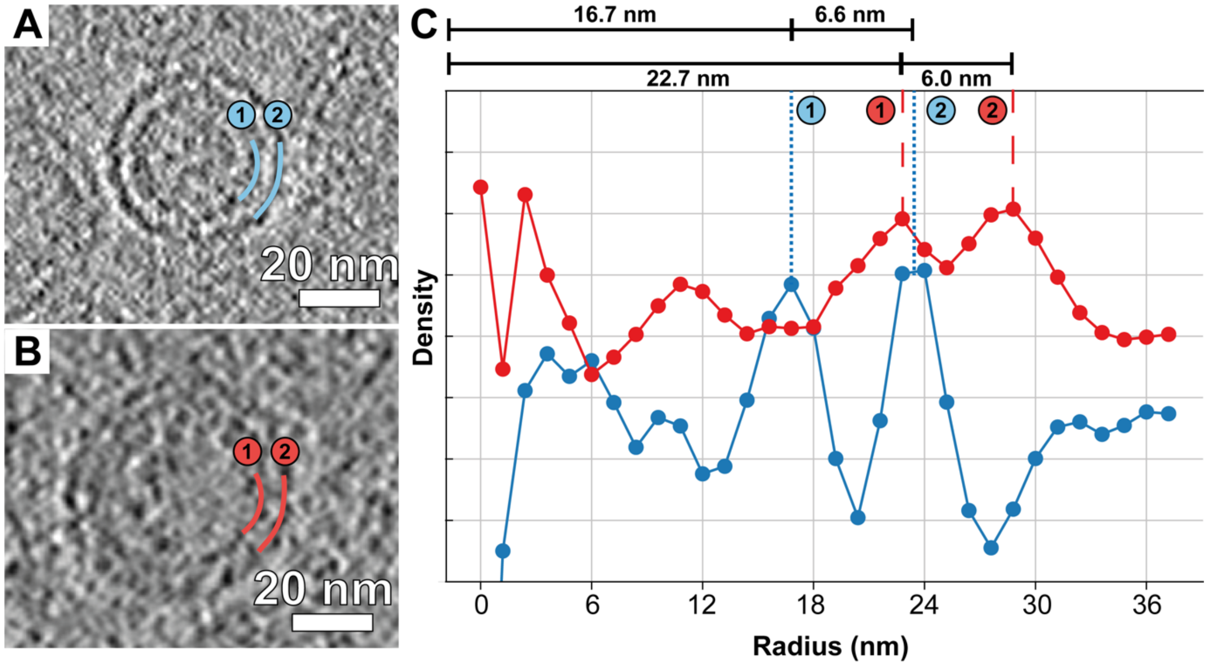


**Fig. S12: Radial plots of VLPs averaged for 5 particles each. A:** Example of a small nuclear VLP and **B:** example of a large cytoplasmic VLP as visualized by tomographic slices. Numbering indicates the two layers formed by Gag and ribonucleocapsid (gRNA+NC) **C:** Radial density plot of the VLPs averaged from 5 particles each. Numbers indicate the density peak locations as shown in A and B. The overall particle diameter determined by radial density average plots is 46.4 nm for the small nuclear and 57.4 nm for the large cytoplasmic VLPs. Pixel size is 11.8 Å at 4x binning.

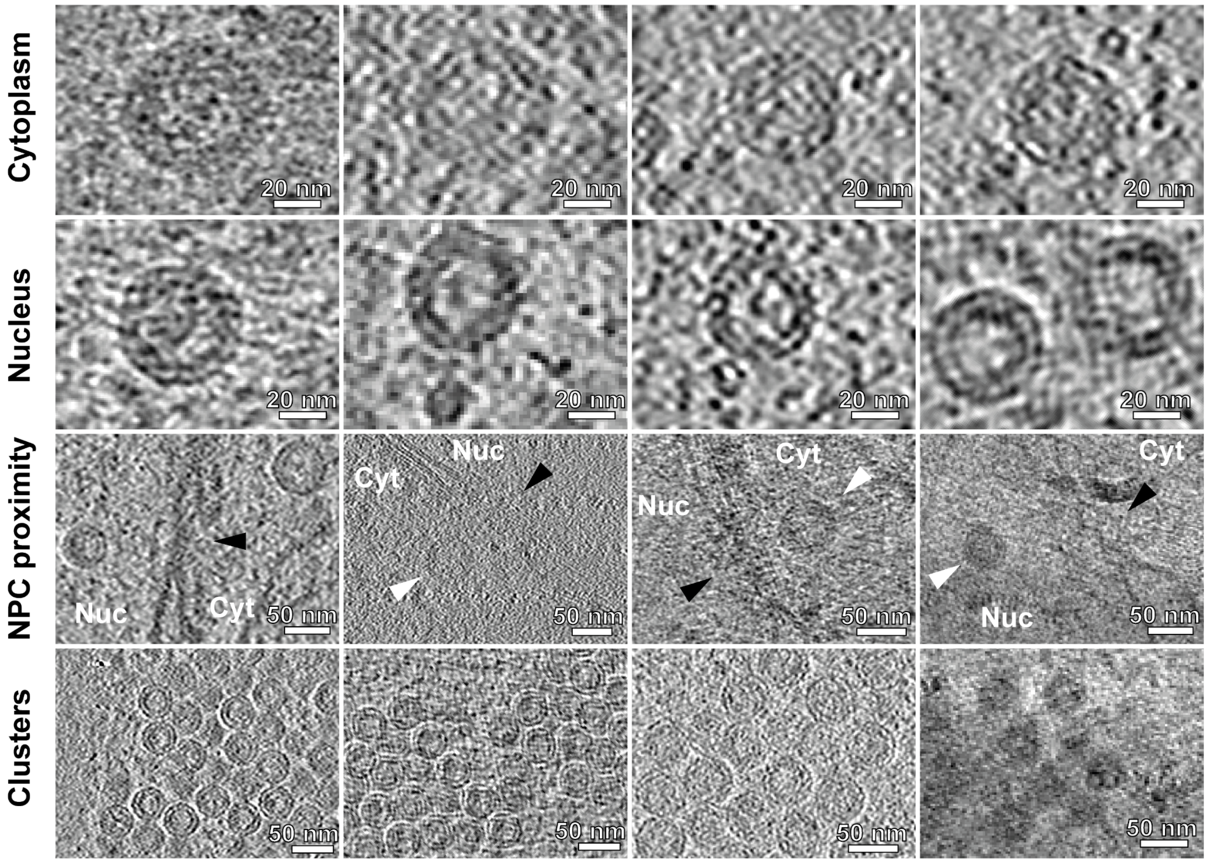


**Fig. S13: Additional examples of in situ observations of *copia* capsids.** Examples for capsids in the cytoplasm (Cytoplasm, Cyt) and nucleus (Nucleus, Nuc) shown to scale for comparison. Additional examples of capsids close to the nuclear pore (NPC proximity) and in nuclear clusters (Clusters). White arrowheads indicate *copia* VLPs, black arrowheads indicate nuclear pores. Genotypes: *tj>EosFP::Sec13*, *LifeAct::GFP*, *GFP::Nup358*

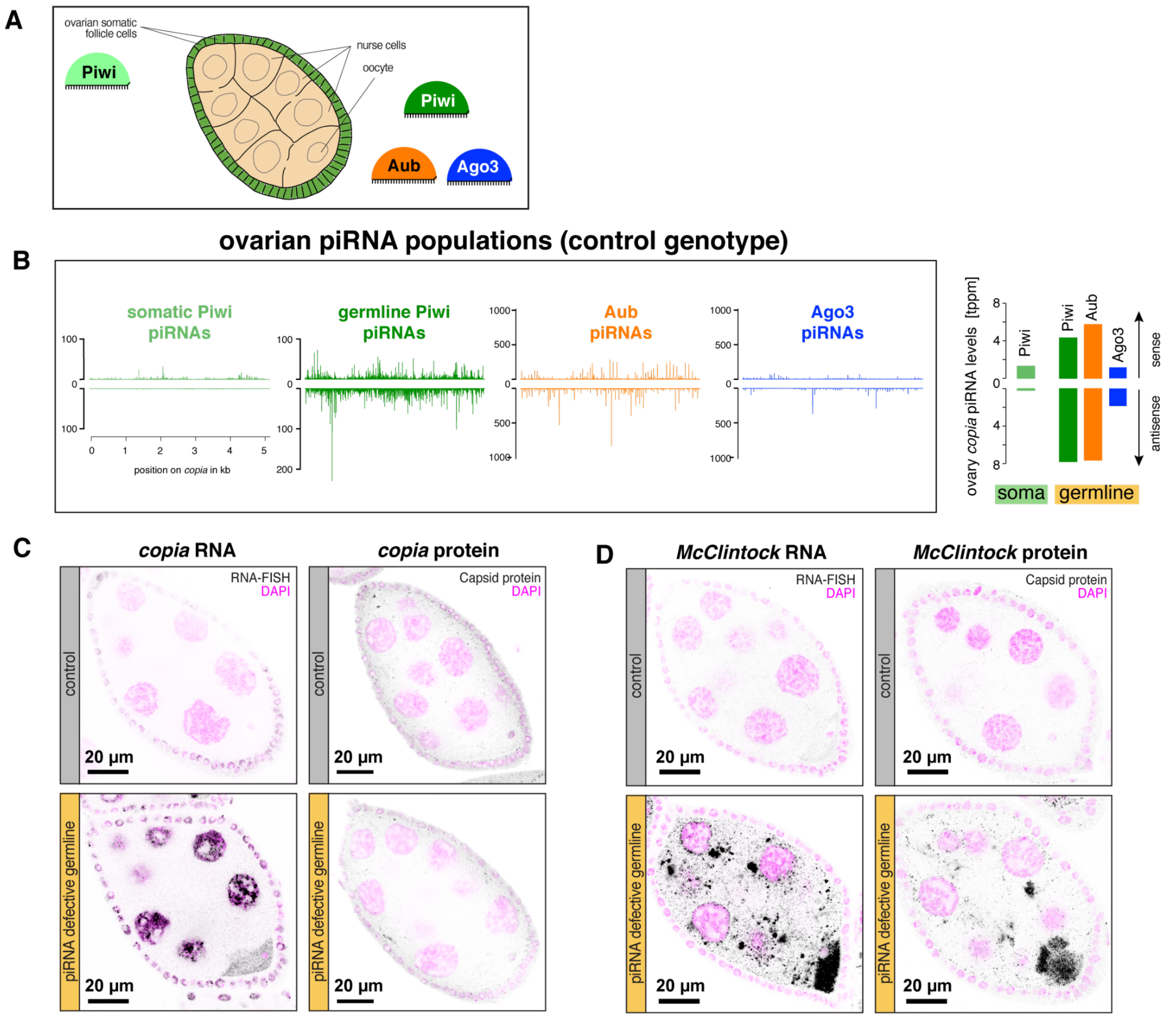


**Fig. S14: Tissue specific ovarian PIWI protein and piRNA populations and levels targeting *copia* in ovaries and comparison of transcript and protein localization for *copia* and *McClintock*. A:** Schematic stage 6/7 egg chamber showing the cell-type specific expression of different PIWI-clade proteins. The ovarian soma only harbors "somaPiwi" (in light green), while the ovarian germline cells express germline Piwi (dark green), Aub (orange), and Ago3 (blue). **B:** Oovarian normalized IP-small RNA-seq data [2] with sense and antisense piRNA profiles along the *copia* consensus sequence for somaPiwi, germlinePiwi, Aub and Ago3 (left panel) and sums of same piRNA populations (right panel, same data as in Figure 5C). Note the profound difference in tissue specific silencing capacity of *copia* in the ovarian germline versus the ovarian soma. **C, D :** Inverted color confocal micrographs showing stage 6/7 egg chambers of germline specific control and piRNA pathway knockdowns (*MTD> white sh* and *MTD>aub ago3 double sh,* respectively) as indicated. **C** shows *copia* smFISH (left panels) and *copia* Gag localisation (right panels) in black and DAPI in magenta. **D** displays *McClintock* smFISH (left panels) and McClintock Gag staining (right panels) in black and DAPI in magenta. Note the strong accumulation of *McClintock* transcripts and Gag in piRNA defective nurse cell cytoplasm and their accumulation in the oocyte in **D**. In contrast, *copia* transcripts are strongly detected in nurse cell nuclei and show only minor oocyte accumulation. We fail to detect any significant *copia* Gag expression or localisation in germline cells of the indicated control or piRNA pathway knockdowns in **C**.

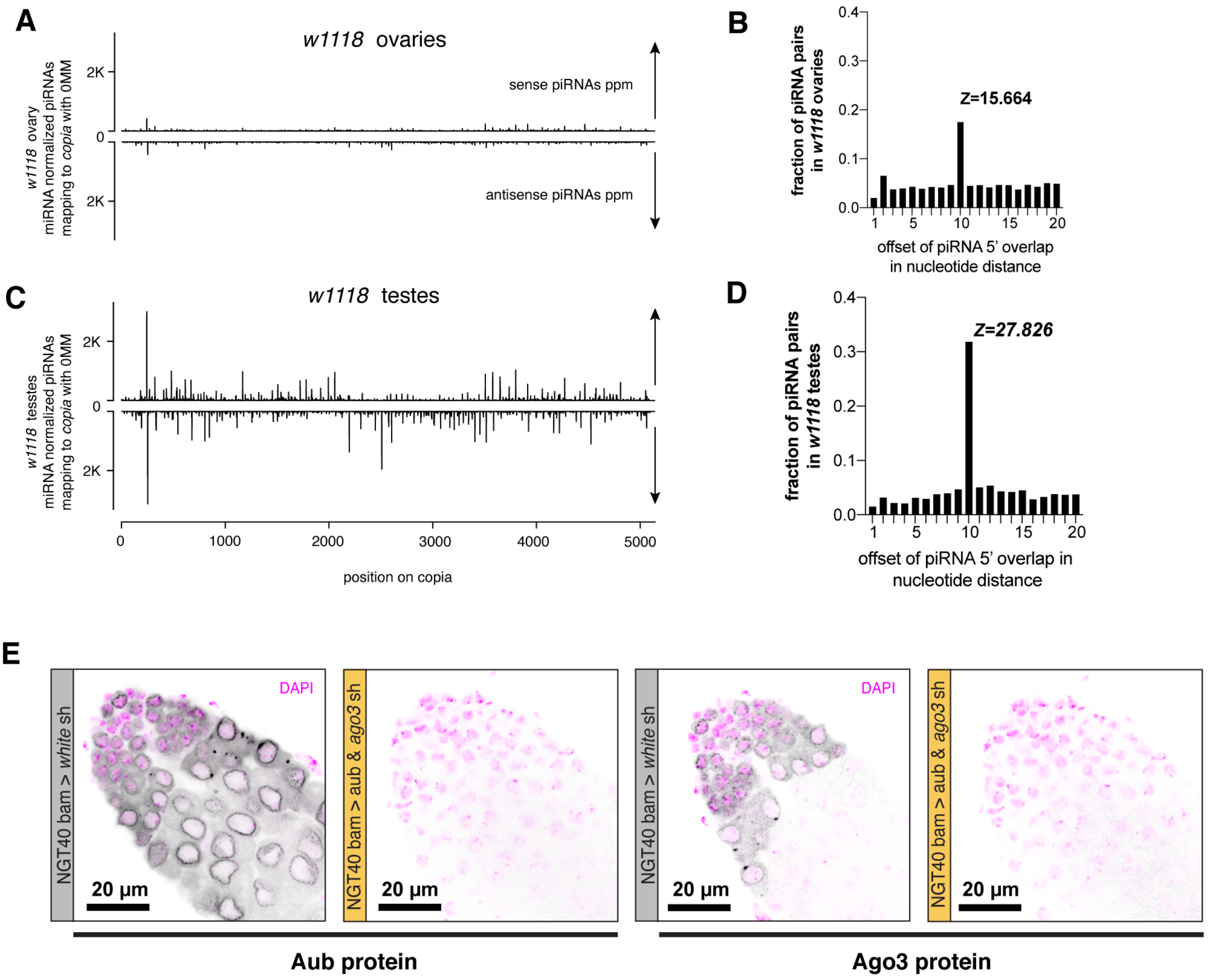


**Fig. S15: Comparison of *copia* regulation in the male and female gonads and validation of the germline specific knockdowns of *aubergine* and *ago3* in testes.** **A-D:** miRNA normalized *copia* piRNA data for *white*$^{1118}$ ovaries **A,B** and testes **C,D**. **A,C** show *copia* piRNAs profiles along the *copia* sequence. **B,D** show the fractions (y axis) of piRNA pairs with N overlaps (x-axis) and the Z score for the 10 base pair piRNA overlaps versus all other piRNA length overlaps plotted. The data shown indicate that the piRNA pathway targets *copia* transcripts in a stronger manner in testes versus ovaries. **E:** Confocal micrographs of testes of germline specific knockdowns of control and aub ago3 double knockdowns (*NGT40&bam>white sh* and *NGT40&bam>aub ago3 sh,* respectively) stained for Aub (left panels) and Ago3 (right panels) (shown in black) and DAPI (magenta). The data in E shows that the experimental depletion of Aub and Ago3 is efficient.
